# The enemy of my enemy: A bacterial antagonist rescues *Chlamydomonas* by inactivating a toxin from a rival

**DOI:** 10.64898/2026.11.05.764488

**Authors:** Trang Vuong, Constanze Schultz, Harikumar R. Suma, Patrick Then, Sankar Jana, Michael Schmitt, Jürgen Popp, Pierre Stallforth, Maria Mittag

**Affiliations:** Matthias Schleiden Institute of Genetics, Bioinformatics and Molecular Botany, Friedrich Schiller University Jena, 07743 Jena, Germany; Leibniz Institute of Photonic Technology (IPHT), Member of Leibniz Health Technologies, Member of the Leibniz Center for Photonics in Infection Research (LPI), Albert-Einstein-Str. 9, 07745 Jena, Germany; Department of Paleobiotechnology, Leibniz Institute for Natural Product Research and Infection Biology, Hans Knöll Institute (HKI), Beutenbergstraße 11a, 07745 Jena, Germany; Cluster of Excellence Balance of the Microverse, Friedrich Schiller University, 07743 Jena, Germany; Institute of Physical Chemistry (IPC) and Abbe Center of Photonics (ACP), Member of the Leibniz Center for Photonics in Infection Research (LPI), Friedrich Schiller University Jena, Helmholtzweg 4, 07743 Jena, Germany; Institute of Organic Chemistry and Macromolecular Chemistry, Friedrich Schiller University Jena, Humboldtstrasse 10, 07743 Jena, Germany

**Author notes:** Author for correspondence: Maria Mittag. Institute for a Sustainable Hydrogen Economy (IHE), Forschungszentrum Jülich GmbH, An der deutschen Welle 7a, 52428 Jülich, Germany. Microbiome Interactions and Engineering Group, Department of Biotechnology and Biomedicine, Technical University of DenmarkSøltofts Plads, 221, 266, 2800 Kgs. Lyngby, Denmark.

## Abstract

Microbial interactions are shaped by multiple community members, yet the underlying mechanisms remain poorly understood. Here, we investigated a tripartite interaction between the microalga *Chlamydomonas reinhardtii* and two *Pseudomonas* bacteria that coexist in the same soil habitat. In pairwise cultures, all three interactions are amensalistic (-like): *Pseudomonas protegens* Pf-5 antagonizes both *C. reinhardtii* and *Pseudomonas fluorescens* HKI0770 without benefit, while *P. fluorescens* inhibits *C. reinhardtii* without profiting from the alga. However, in a tripartite culture, *P. fluorescens* protects *C. reinhardtii* from *P. protegens* and promotes algal recovery. This protection is not stopped when the used carbon source, acetate, is supplemented. The *P. fluorescens* rescue effect is associated with reduced levels of the polyyne toxin protegencin produced by *P. protegens*. Protegencin levels, detected by Raman microspectroscopy, are decreased by application of spent medium or protein extracts from *P. fluorescens*, while heat- and protease-treatment abolish this activity. A heat-sensitive, proteinaceous factor, likely an enzyme(s), appears to contribute to protegencin inactivation. Our results show that *P. fluorescens* rescues *C. reinhardtii* by inactivating protegencin, despite being inhibited by *P. protegens*. These findings demonstrate the importance of investigating higher-order effects, since additional community members can transform pairwise antagonistic interactions into conditional protection through toxin inactivation.

## Introduction

Microorganisms rarely exist as isolated populations but instead form complex communities in which interactions among multiple species shape community structure and ecological function^1^. These interactions range from cooperation and resource exchange to competition and antagonism and are often mediated by nutrient availability, signaling molecules, and specialized metabolites^2^. Among different types of microbial relationships, amensalism represents an important but relatively understudied form of microbial interaction^1^. In this interaction, one partner is negatively affected while the other experiences no detectable effect. However, the outcome of such pairwise interactions can be strongly altered by the presence of additional community members, leading to emergent effects that cannot be predicted from individual interactions alone^3^.

Microalgae are key primary producers in aquatic and terrestrial ecosystems, forming the basis of many food webs and contributing substantially to global CO_2_ fixation through photosynthesis^4–6^. Microalgal growth, physiology, and survival are influenced by abiotic conditions but also by interactions with other microorganisms, particularly bacteria^7^. Algal-bacterial associations can affect nutrient availability, metabolic activity, and stress responses, thereby shaping the ecological success of both partners^8–10^. Many of these relationships are mediated by exchanges of chemical mediators within the phycosphere, the microenvironment surrounding algal cells, where algae and bacteria exchange metabolites that can influence each other’s physiology and growth^11–13^. Beyond their prominent ecological consequences, biotic interactions have even contributed to algal evolution and diversification^14^. Understanding these relationships is therefore essential for elucidating how microbial communities influence algal performance and how biotic associations contribute to algal diversification and adaptation.

In recent years, the unicellular green alga *Chlamydomonas reinhardtii* has emerged as a model for studying algal–bacterial interactions^9,15^. Originally isolated from potato soil^16^, *C. reinhardtii* and closely related *Chlamydomonas* strains have also been repeatedly detected in diverse soil environments^17^, including rice fields^18–20^, where it coexists in complex microbial communities. Rather than interacting randomly with surrounding microorganisms, *C. reinhardtii* assembles a characteristic microbiota whose composition shares remarkable taxonomic and functional similarities with the root microbiota of higher plants, suggesting that conserved mechanisms govern host–microbe interactions across the green lineage^21^. The underlying bacterial community is dominated by members of the Pseudomonadota (formerly called Proteobacteria), suggesting that interactions with *Pseudomonas* species are a common feature of the algal natural ecology. Among *Pseudomonas* bacteria, *Pseudomonas protegens* Pf-5 (hereafter named *P. protegens*) represents a prominent antagonist of *C. reinhardtii*^22^. This bacterium occurs in soil-associated environments, including potato soil and rice fields^23,24^. It produces two major toxins, orfamide A and the polyyne toxin, protegencin, which target *C. reinhardtii* and act on different timescales. Orfamide A impairs algal physiology by causing flagellar loss, whereas protegencin destroys the eyespot and induces rapid cell lysis^22,25,26^. In addition to orfamide A and protegencin, *P. protegens* produces a diverse arsenal of secondary metabolites with antialgal activities, including rhizoxin S2, pyrrolnitrin, pyoluteorin, and 2,4-diacetylphloroglucinol (DAPG)^27^.

Here, we investigated how a third bacterial partner, *Pseudomonas fluorescens* HKI0770 (see Methods; hereafter named *P. fluorescens*)^28^, a species which is also present in rice fields^29,30^ may influence the antagonistic interaction between *C. reinhardtii* and *P. protegens*. We identified *P. fluorescens* as an amensalist of *C. reinhardtii* since it inhibits algal growth without benefiting from the alga. Furthermore, *P. fluorescens* acts as a conditional protector that rescues *C. reinhardtii* from *P. protegens*-mediated toxicity despite itself being inhibited by *P. protegens*. We show that this protective effect is independent of acetate, the provided carbon source for all three microorganisms. Using Raman microspectroscopy, a label-free approach that enables detection of chemical signatures at the single-cell level, we show that the rescue mechanism involves a reduction in the toxin protegencin released by *P. protegens* within the tripartite consortium. We provide evidence that at least one *P. fluorescens*-derived enzyme contributes to toxin inactivation. Our findings reveal how multispecies microbial interactions can shift antagonistic relationships toward protective outcomes and highlight the importance of community context in understanding microbial warfare.

## Results and Discussion

### Both *P. protegens* and *P. fluorescens* engage in amensalistic interactions with *C. reinhardtii*

The unicellular green alga *C. reinhardtii* and its antagonistic bacterium *P. protegens* are both present in rice field soil. We wondered whether another rice soil bacterium, *P. fluorescens*^28–30^ may exert an inhibitory effect on *C. reinhardtii*. We therefore investigated the interaction between *C. reinhardtii* and *P. fluorescens*, in comparison to the algal interaction with *P. protegens*. We grew the cultures in Tris-Acetate-Phosphate (TAP) medium, in which *C. reinhardtii* grows mixotrophically (Harris, 1989). Acetate is present in rice field soils^31,32^, and serves as a carbon source for the growth of all three microorganisms^27,33,34^. In coculture with *P. fluorescens*, algal growth seemed reduced, as indicated by the lack of green color (days 1 and 2) or lighter green coloration (days 3 to 10) due to photosynthetic algal pigments, compared with a *C. reinhardtii* monoculture. However, in contrast to the interaction of *C. reinhardtii* and *P. protegens*, which fully bleached the algal cultures over a longer time frame (days 1 to 7), *P. fluorescens* had a weaker but well-visible antagonistic effect (Fig. 1A, S1A).

**Fig. 1.**
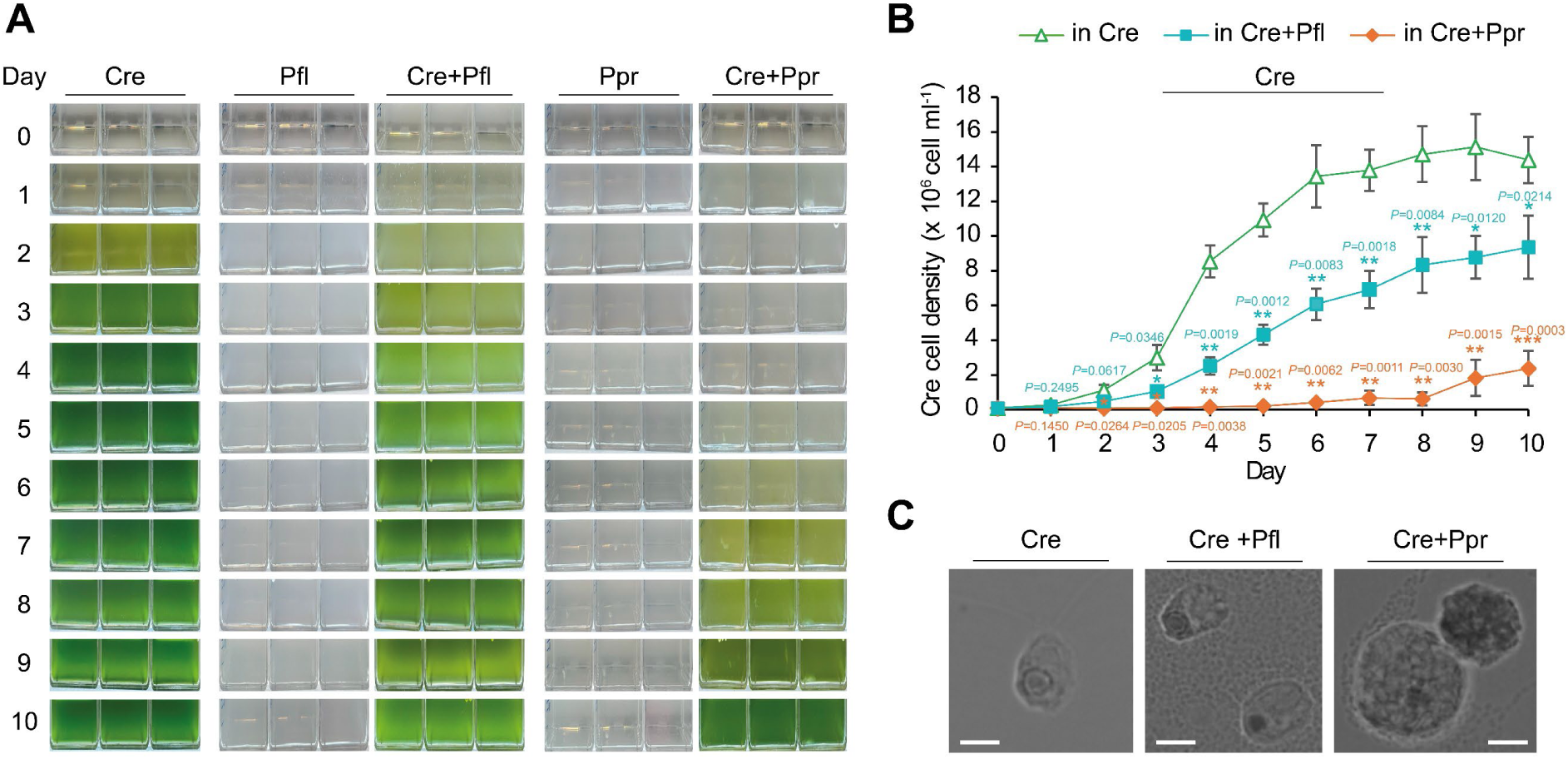
***P. protegens* (Ppr) and *P. fluorescens* (Pfl) exert distinct inhibitory effects on *C. reinhardtii* (Cre). A** Algal and bacterial monocultures and bipartite cultures. The organisms were cultivated in TAP medium at a ratio of 1:250 (Cre:Ppr or Cre:Pfl). Equally treated monocultures of Cre, Pfl and Ppr served as controls. Photographs were taken every day for 10 days. Experiments were repeated three times independently (n=3). Independent replicates 2 and 3 are shown in Fig. S1. **B** The growth of Cre in bipartite cultures compared with its monoculture. Cre cell numbers were determined using a Neubauer improved chamber (see Methods). Values represent the mean of three independent biological replicates (n=3). Error bars indicate the standard deviation (SD). Statistical significance was determined by comparing cell numbers in each condition with those in the Cre monoculture using an unpaired two-tailed Welch’s *t*-test (unequal variances). Exact *P*-values, *t*-values and degrees of freedom are reported in Table S1. Significance levels are indicated as follows: \**P* < 0.05, \*\**P* < 0.01 and \*\*\**P* < 0.001; unlabeled comparisons are not statistically significant. **C** Bright-field images of Cre cells in cocultures with *P. fluorescens* and *P. pseudomonas* bacteria. Scale bars = 5 µm. Imaging was performed using a spinning disk confocal laser scanning microscope (AxioObserver.Z1/7, Carl Zeiss) equipped with a 100x/1.40 NA Plan-Apochromat oil-immersion DIC M27 objective lens (Carl Zeiss). Representative images from two independent experiments (n=2) are shown.

We next studied all bipartite interactions of the three microorganisms by monitoring the growth of all partners in their respective cocultures. While the algal growth is significantly reduced by *P. fluorescens* compared to its monoculture, *P. protegens* causes even a stronger decline in algal cell density, which remained close to zero until day 8 (Fig. 1B, Table S1). Brightfield microscopy images of single algal cells in bacterial cocultures confirmed that *P. protegens* alters the algal usual oval shape^22^, resulting in a circle-like shape with granular structures inside the cell. *P. fluorescens* seems not to cause any morphological changes (Fig. 1C).

We also monitored the growth of both bacteria in coculture with *C. reinhardtii*. Although both bacteria tended to reach higher cell densities in coculture than in their respective monocultures, the differences were not statistically significant (Fig. S1B, C; Table S2). As both bacteria negatively affected the alga in its growth, while neither bacterium benefited significantly from *C. reinhardtii*, amensalistic-like interactions^1^ are indicated in the used TAP medium. It has been shown before that *P. protegens* can acquire micronutrients from *C. reinhardtii* under nutrient-limiting conditions, in which trace elements were omitted from the TAP medium used in our experiments^22^ . This may also explain the tendency of the bacteria to reach higher cell densities in coculture even though it is not significant; they might uptake slightly higher amounts of micronutrients in independent biological experiments.

### *P. protegens* exerts an amensalistic effect on *P. fluorescens*

Next, we examined the interaction between the two bacteria in coculture. *P. protegens* cell numbers in coculture with *P. fluorescens* were not significantly different from those in its monoculture (Fig. 2A; Table S3). However, *P. protegens* strongly suppressed *P. fluorescens* in bacterial coculture, resulting in substantially lower cell numbers of *P. fluorescens* than in its monoculture (Fig. 2B, Table S3). Therefore, *P. protegens* behaves as an amensalist towards *P. fluorescens*^1^. *P. protegens* is known to produce metabolites having antimicrobial activities, such as 2,4-diacetylphloroglucinol (DAPG)^35^, pyoluteorin^36^, pyrrolnitrin^37^ and protegencin^38^. These metabolites may likely contribute to the growth inhibition effects of *P. protegens* on *P. fluorescens*.

**Fig. 2.**
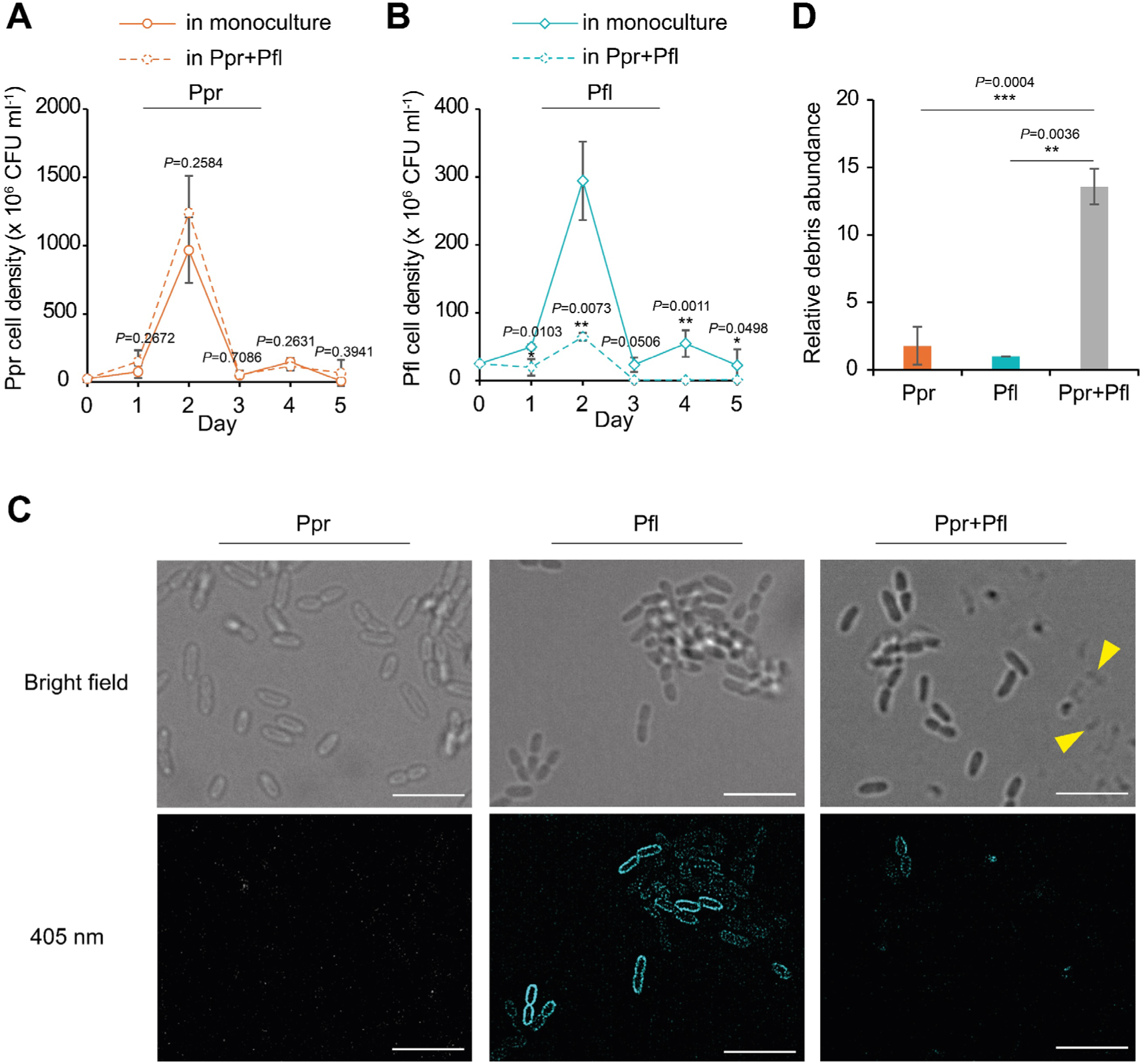
Amensalistic interaction between *P. protegens* (Ppr) and *P. fluorescens* (Pfl). **A** The growth of Ppr in coculture with Pfl compared to its monoculture over 5 days. **B** The growth of Pfl in coculture with Ppr compared to its monoculture over 5 days. **A, B** Bacterial cell numbers were determined by colony-forming unit (CFU) assays (Fig. S2, see Methods). **C** Representative brightfield and fluorescence images of Ppr, Pfl, and their coculture (Ppr+Pfl) on day 2. Images were taken using an Elyra Zeiss microscope, with fluorescence acquired in Structured Illumination Microscopy (SIM) mode (see Methods). Pfl was identified by its intrinsic autofluorescence upon 405 nm excitation (turquoise channel), whereas Ppr showed only very low residual fluorescence under these conditions. Yellow arrows indicate cell lysis of Pfl. Scale bars = 5 µm. Representative images from three independent experiments (n=3) are shown. Additional images of the bacterial interactions and cell debris are shown in Fig. S3, respectively. **D** Relative debris abundance in monocultures of Ppr and Pfl and in the Ppr+Pfl coculture. The debris abundance in the Pfl monoculture was set to 1, and values for Ppr monoculture and Ppr+Pfl coculture were normalized accordingly. **A, B, D** Data are presented as the mean of three independent biological replicates (n=3). Error bars represent the standard deviation (SD). Statistical significance was assessed using an unpaired two-tailed Welch’s *t*-test (unequal variances). Significance levels are indicated as follows: \**P* < 0.05, \*\**P* < 0.01 and \*\*\**P* < 0.001; unlabeled comparisons are not statistically significant. Exact *P*-values, *t*-values and degrees of freedom are reported in Tables S3 and S4, respectively.

The interaction between the two bacteria was further examined by fluorescence microscopy. *P. fluorescens* exhibits intrinsic autofluorescence under 405 nm excitation, whereas *P. protegens* shows only very low residual levels of fluorescence under these conditions, allowing the two bacterial populations to be distinguished in coculture. Consistent with the growth data, only a few *P. fluorescens* cells were observed in the coculture with *P. protegens* compared to its monoculture (Fig. 2C, S3A). Moreover, the background in monocultures was largely free of particulate material in corresponding brightfield images. In contrast, the bacterial coculture contained cellular debris (Fig. 2C, yellow arrows; Fig. S3A, B) and the number of debris was significantly higher than in the monocultures (Fig. 2D, Table S4). The increased abundance of debris, together with the strongly reduced autofluorescence of *P. fluorescens* in the coculture, supports the hypothesis that *P. protegens* induces lysis of *P. fluorescens* cells. This may involve a mechanism similar to that involved in algal lysis caused by the above-mentioned polyyne, protegencin (Hotter et al., 2021). Given its broad antimicrobial activity against diverse bacteria^38^, protegencin may lead to *P. fluorescens* cell lysis.

### *P. fluorescens* acts as a conditional protector, rescuing *C. reinhardtii* only in the presence of *P. protegens*

We next investigated the combined effects of the two amensalistic bacterial partners on *C. reinhardtii* and their interactions within a tripartite community. Strikingly, in the presence of both bacteria, *C. reinhardtii* started to recover from day 2 onward, as indicated by the restoration of green pigmentation. In contrast, this recovery was not observed in the algal coculture with *P. protegens*. The timing of recovery (magenta arrow) varied slightly between independent biological replicates, which are therefore shown separately (Fig. 3A, 4SA, C). Although *P. fluorescens* inhibited *C. reinhardtii* growth in the bipartite culture, it markedly protected the alga from the antagonistic activity of *P. protegens* in the tripartite culture.

**Fig. 3.**
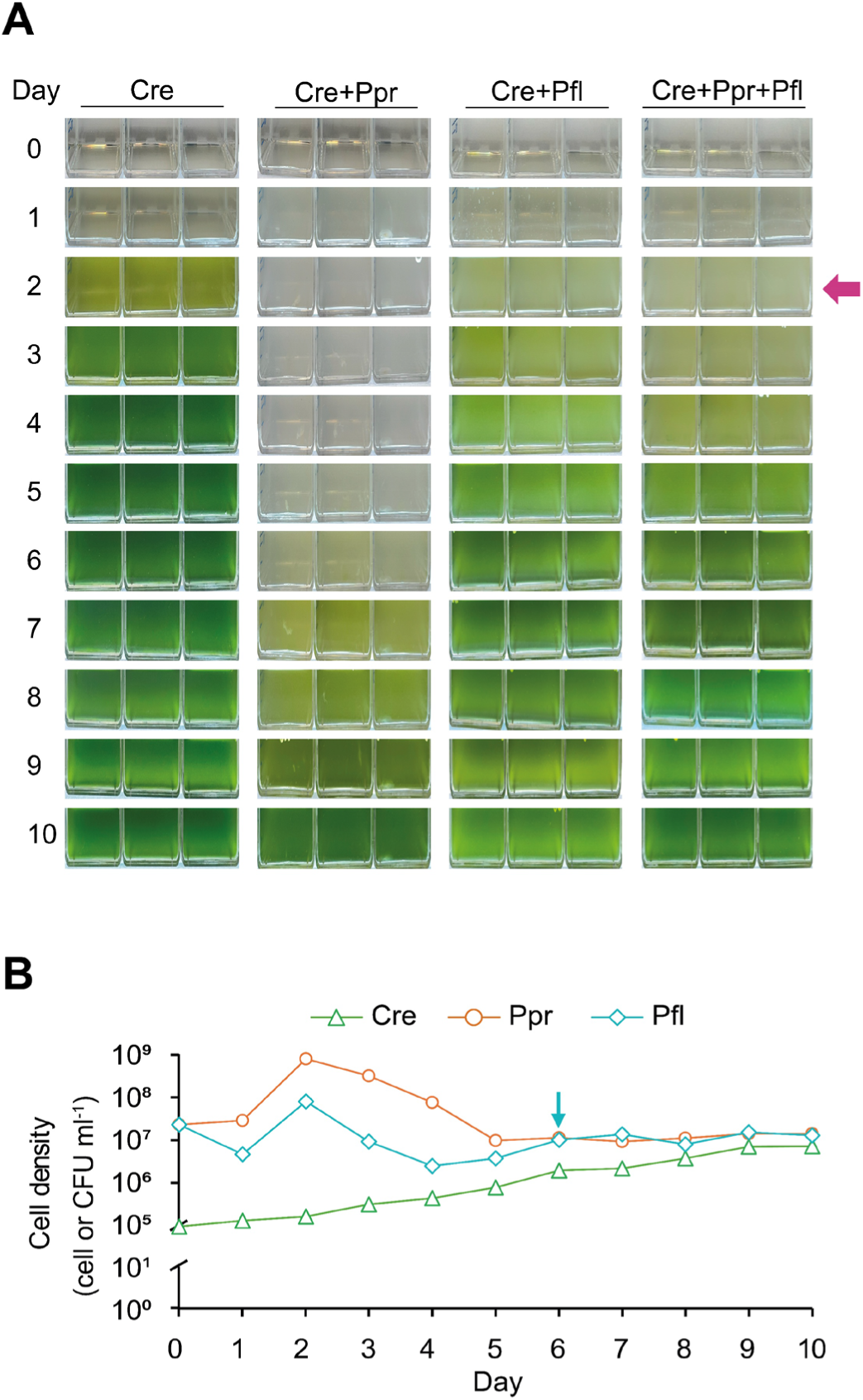
The amensalist *P. fluorescens* (Pfl) becomes a conditional protector, rescuing *C. reinhardtii* (Cre) from *P. protegens* (Ppr) in a tripartite culture. **A** Mono-, bi- and tripartite cultures of Cre, Ppr and Pfl. Photographs were taken every day for 10 days. The magenta arrow indicates recovery of Cre. Due to variation in Cre and Pfl recovery among replicates, individual replicates are shown. **B** The growth of Cre, Ppr and Pfl in a tripartite culture. Bacterial cell numbers were determined by colony-forming unit (CFU) assays. Cre cell numbers were determined using a Neubauer improved chamber (see Methods). The turquoise arrow indicates the recovery of Pfl. **A-B** The experiment was repeated three times independently (n=3). Independent replicates 2 and 3 are shown in Fig. S4.

**Fig. 4.**
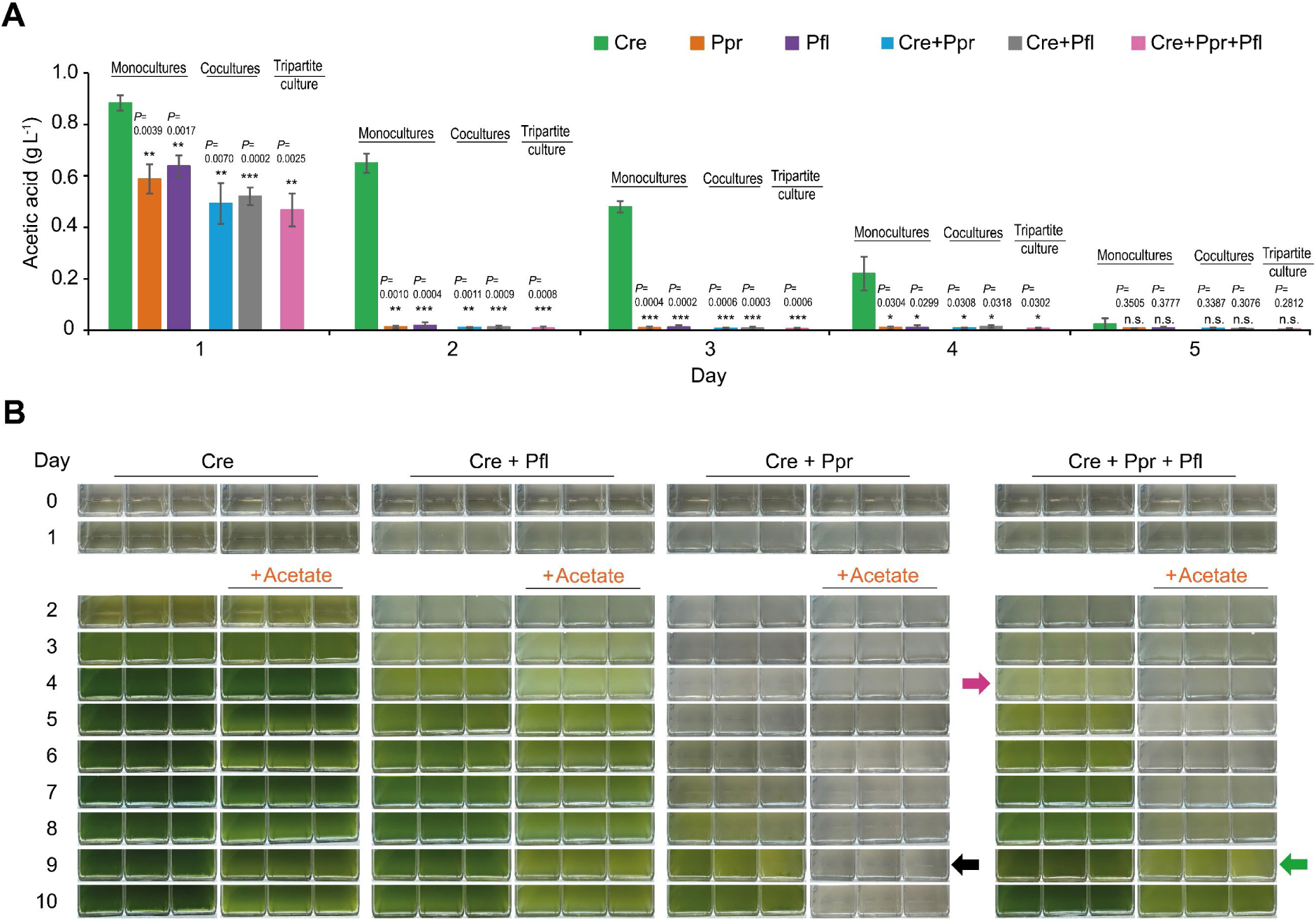
The availability of acetate does not stop the rescue function of *P. fluorescens*. **A** The levels of acetate in the mono-, bi- and tripartite cultures of *C. reinhardtii* (Cre), *P. protegens* (Ppr) and *P. fluorescens* (Pfl) over 5 days. Statistical significance was determined using an unpaired Student’s *t*-test (Welch’s test). Values were normalized to the acetate levels in Cre monoculture. The experiments were performed with three independent biological replicates (n=3), each consisting of three technical replicates. Error bars represent the standard deviation (SD) of the biological replicates. Statistical significance was determined using an unpaired two-tailed Welch’s *t*-test (unequal variances). Significance levels are indicated as follows: \**P* < 0.05, \*\**P* < 0.01 and \*\*\**P* < 0.001. Exact *P*-values, *t*-values and degrees of freedom are reported in Table S5, respectively. **B** Acetate independence assay in the tripartite culture of Cre, Ppr and Pfl. Monocultures of Cre and cocultures of Cre+Ppr and Cre+Pfl were included as controls. For each condition, six flasks were inoculated. On day 2, acetic acid buffered to pH 7 was added to three flasks in the same concentration as in TAP medium, while the remaining three flasks were left untreated. Magenta and green arrows indicate recovery of Cre, whereas the black arrow indicates bleaching of the Cre+Ppr coculture. **A-B** All cultures were started in TAP medium at the initial ratio of 1:250:250 (Cre:Pfl:Ppr). The experiments were repeated three times independently (n=3). Independent biological replicates 2 and 3 are shown in Fig. S5.

To further study the dynamics in the tripartite culture, we monitored the growth of each microorganism. Over 10 days, *C. reinhardtii* gradually recovered from 10^5^ cell ml^-1^ to 7.7×10^6^ cell ml^-1^. *P. protegens* suppressed *P. fluorescens* during the first four days. After day 2, both bacterial populations declined whereby *P. fluorescens* was stronger reduced. Interestingly, from days 6 to 7, *P. fluorescens* began to recover its growth slightly from 4×10^6^ cell ml^-1^ to 1.7×10^7^ cell ml^-1^ (Fig. 3B). The growth curves of the three partners in independent biological replicates 2 and 3 showed similar results (Fig. S4B and D). The recovery of *P. fluorescens* may be associated with the concurrent recovery of *C. reinhardtii*, potentially reflecting improved resource availability or changes in the cocultural environment. *C. reinhardtii* is known to release a range of metabolites, including certain B vitamins and methionine^23^ and *P. fluorescens* has been reported to use methionine as a carbon and nitrogen source^39^. Thus, algal-derived metabolites could potentially contribute to the recovery of *P. fluorescens* in the tripartite culture (Fig. 3B, 4SB, D).

Together, these results reveal an unexpected role of *P. fluorescens*: despite being inhibited by *P. protegens* and exerting an antagonistic effect on *C. reinhardtii* in respective bipartite cultures, it functions as a conditional protector of the alga in the presence of the strong amensalist.

### The algal rescue effect by *P. fluorescens* is not stopped when acetate is supplemented

In complex microbial communities, microorganisms may compete for the carbon source, which could influence their growth and interactions. To determine whether *P. fluorescens*-mediated rescue of *C. reinhardtii* depends on the carbon source used by all three partners, we investigated acetate availability. We monitored acetate levels in mono-, bi-, and tripartite cultures of *C. reinhardtii*, *P. protegens*, and *P. fluorescens* over five days. In the *C. reinhardtii* monoculture, acetate levels gradually decreased and were nearly depleted by day 5 (Fig. 4A, Table S5). In contrast, in all cultures containing bacteria, including the bacterial monoculture, bipartite, and tripartite cultures, acetate was already largely depleted by day 2 (Fig. 4A, Table S5). As acetate becomes depleted and *C. reinhardtii* gradually recovers, the bacteria, particularly *P. fluorescens*, may rely on algal-derived nutrients, supporting the possibility of metabolic exchange between the partners, as mentioned before.

Next, we tested whether *P. fluorescens* protection depends on acetate availability. For each culture condition, culture flasks were prepared in duplicates. On day 2, when acetate was almost completely depleted (Fig. 4A), acetate was added to one set of flasks, while the other set of flasks was maintained without acetate addition as controls. The amount of added acetate, buffered to pH 7, was the same as in TAP medium, thus avoiding potential side effects of acetate. Following acetate supplementation, both *P. fluorescens* and *P. protegens* exhibited enhanced antagonistic effects toward *C. reinhardtii* in their respective bipartite cultures (Fig. 4B, S5). In the *C. reinhardtii*–*P. protegens* coculture, acetate supplementation prevented algal recovery throughout the 10-day growth period (Fig. 4B). The control, i.e. the tripartite cultures, in which acetate was not added, began to recover on day 4 (magenta arrow). Acetate supplementation still allowed algal recovery in the tripartite cultures, but with a delay of some days (Fig. 4B). Thus, the algae recovered by day 9 (green arrow), despite the addition of acetate. Although acetate supplementation prolonged *P. protegens* antagonism, algal recovery eventually occurred in the presence of *P. fluorescens* (green arrow) compared to the bipartite culture of *C. reinhardtii* and *P. protegens* (black arrow). Similar data were obtained in two other independent biological replicates (Fig. S5). These data suggest that acetate availability contributes to the protective effect of *P. fluorescens*, but additional mechanisms may also play a role.

### *P. fluorescens* reduces the level of the protegencin toxin secreted by *P. protegens*

The underlying tripartite interaction mechanism was further investigated using fluorescence *in situ* hybridization (FISH). In the coculture of *C. reinhardtii* and *P. fluorescens*, algal cells maintained their oval shape as observed in monoculture (Fig. 5A, first row; Fig. S6) and shown before (Fig. 1C). Bacterial cells densely surrounded the algal cells (Fig. 5A, second row; Fig. S6). In the coculture with *P. protegens*, the algae were also surrounded by bacteria but appeared severely damaged and lysed (Fig. 5A, third row; Fig. S6). The lysis effect is largely due to the major polyyne toxin, protegencin, produced by *P. protegens*^25^. In the tripartite culture, both lysed and intact algal cells were observed (Fig. 5A, last row, yellow arrows; Fig. S6), suggesting that protegencin-mediated lytic activity may be diminished in the presence of *P. fluorescens*. Together, these observations support a protective role of *P. fluorescens* against *P. protegens* toxicity on *C. reinhardtii* and prompted us to examine the core mechanism further.

**Fig. 5.**
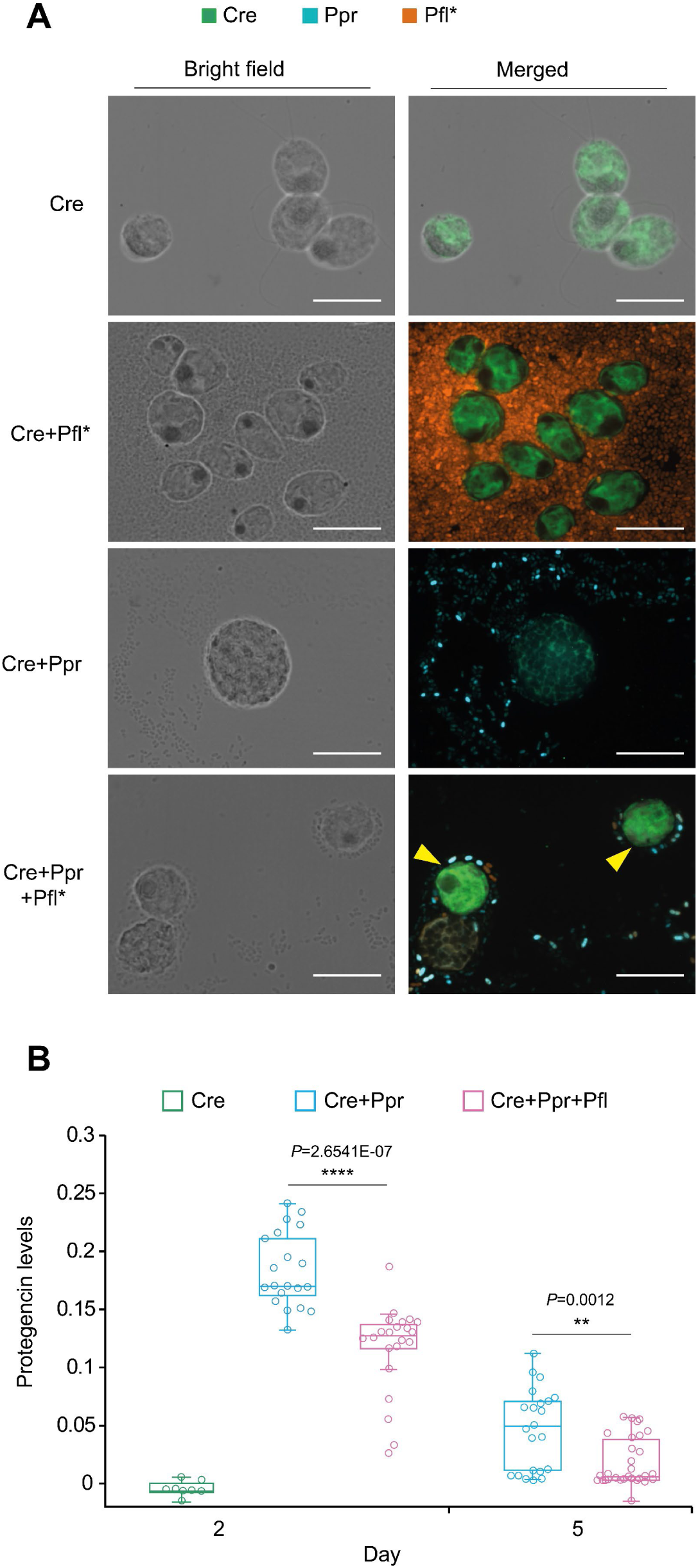
***P. fluorescens-*mediated protection of *C. reinhardtii* is associated with reduced protegencin levels. A** FISH images visualizing the bi- and tripartite interactions. Imaging was performed using a spinning disk confocal laser scanning microscope (AxioObserver.Z1/7, Carl Zeiss) equipped with a 100x/1.40 NA Plan-Apochromat oil-immersion DIC M27 objective lens (Carl Zeiss) (see Methods). Merged images were generated using the 488 nm channel for *C. reinhardtii* (Cre), the 561 nm channel for fluorescent labelled *P. fluorescens*\* (Pfl*), and the 638 nm channel for *P. protegens* (Ppr). An overview of the images acquired at the different wavelengths corresponding to each fluorescent probe is provided in Fig. S6. Bright-field and merged images are shown to provide a clearer view of the interactions among the three organisms. Green channel = Cre, orange channel = Pfl*, blue channel = Ppr. Yellow arrows indicate intact, live Cre cells. Scale bars = 10 µm. Representative images from two independent experiments (n=2) are shown. **B** Protegencin levels were quantified on days 2 and 5 by Raman microspectroscopy using the ratio of the triple-bond signal at (2170 ± 100) cm⁻¹ to the reference signal at (1635 ± 80) cm⁻¹. All cultures were started in TAP medium at the initial ratio of 1:250:250 (Cre: Pfl: Ppr). On day 2, 20 independent cells were measured for the Cre+Ppr coculture and 22 independent cells for the Cre+Ppr+Pfl tripartite culture. On day 5, 22 and 29 independent cells were measured, respectively. A monoculture of Cre on day 2 was included as further control; for this sample, 8 independent cells were measured. The data are presented as box-and-whisker plots, with the center line indicating the median, the box representing the interquartile range (IQR), and the whiskers extending to the minimum and maximum values. Individual values are shown as scattered points, including both inner and outlier points. Statistical significance was determined using an unpaired two-tailed Welch’s *t*-test (unequal variances). Significance levels are indicated as follows: \*\**P* < 0.01, and \*\*\*\**P* < 0.0001. Exact *P*-values, *t*-values and degrees of freedom are reported in Table S6. Values from a second independent dataset, together with additional measurements from days 3 and 4, are shown in Fig. S7.

We next quantified protegencin levels within the algal cells using Raman microspectroscopy developed to examine the algal cells^25^. Raman microspectroscopy is one of the unique methods to quantify polyynes *in situ* in algal cells, as polyynes with their triple bonds, appear in the silent region of the Raman spectrum. In the algal monoculture (day 2), protegencin levels are close to zero (Fig. 5B), while they were strongly increased in bipartite cultures of *C. reinhardtii* and *P. protegens*, especially on day 2 when the bacterial growth is in the highest range (Fig. S1B) as well as on day 5, where bacterial growth declined again. Strikingly, protegencin levels were consistently lower in the tripartite cultures together with *P. fluorescens* at days 2 and 5, compared to cocultures of *C. reinhardtii* and *P. protegens* (Fig. 5B, Table S6). Measurements from an independent dataset showed a similar trend, with a strong reduction on day 5 (Fig. S7A, Table S7A). Additional measurements on days 3 and 4 corroborated these results (Fig. S7B, Table S7B). As the Raman microspectroscopy measurements can specifically detect the carbon-carbon triple bonds of the toxin, our findings strongly suggest that protegencin, which bears four triple bonds, is degraded to a significant extent in the presence of *P. fluorescens*.

### Enzyme(s) from *P. fluorescens* seem to mediate the protegencin degradation

To characterize the *P. fluorescens*-derived protective factor, we investigated at first whether the protective activity may be present in the culture spent medium pointing to its secretion. Spent medium from the tripartite culture was used to cultivate *C. reinhardtii* in mono-or in coculture with *P. protegens* (Methods, Fig. 6A, S8). As acetate was already depleted by day 2 (Fig. 4A), we supplemented the spent medium with nutrients from TAP medium, including acetate (see Methods), to ensure the growth of the organisms. By day 7, *C. reinhardtii* recovered in the coculture grown in the presence of spent medium from *P. fluorescens*, whereas no recovery was observed in the corresponding coculture grown without spent medium (Fig. 6A, S8). This indicates that the protective activity involves factor(s) released into the extracellular environment. We postulated that an enzyme that cleaves the toxin may be responsible for its degradation. Although no such enzyme has yet been described for protegencin, an example of an enzyme modifying a carbon–carbon triple bond by catalyzing the hydration of acetylene to acetaldehyde is known, with an acetylene hydratase^40^. To further investigate whether an enzyme or alternatively a heat-sensitive compound may be involved, the *P. fluorescens* spent medium was heated at 121 °C by autoclaving (see Methods) before inoculation. Heat treatment abolished the protective activity; the coculture remained bleached throughout the experiment (Fig. 6A, S8).

**Fig. 6.**
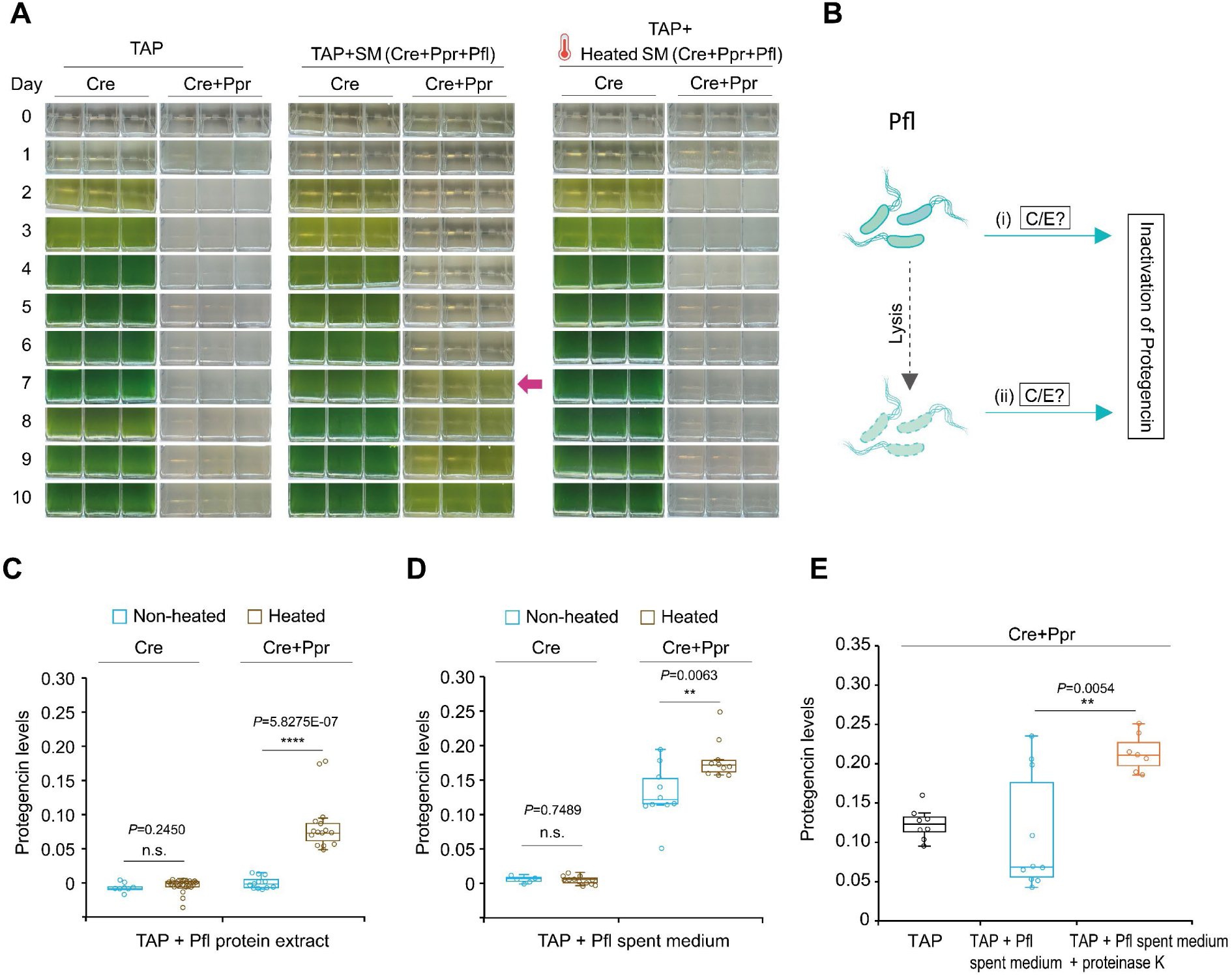
At least one protein, likely an enzyme, from *P. fluorescens* (Pfl) is involved in the algal rescue mechanism. **A** Cocultures of *C. reinhardtii* (Cre) and *P. protegens* (Ppr) in non-heated and heated spent medium (SM) of a tripartite culture [SM (Cre+Ppr+Pfl)]. The magenta arrow indicates the recovery of Cre in non-heated SM. This experiment was repeated 3 times (n=3). The other 2 independent replicates are shown in Fig. S8. **B** Schematic illustration of the two proposed hypotheses for the origin of the Pfl-derived protective factor(s): (i) Pfl actively secretes compound(s) or enzyme(s) into the extracellular medium, or (ii) Pfl cells are lysed, releasing intracellular compound(s) or enzyme(s) that inactivate protegencin. C, compound; E, enzyme. **C** Levels of protegencin in the cocultures of Cre+Ppr in non-heated and heated protein extract of Pfl. Under non-heated treatments, 8 independent cells were measured for Cre monoculture and 12 for Cre+Ppr coculture. Under heated treatments, 28 and 15 independent cells, respectively, were measured. **D** The levels of protegencin in the cocultures of Cre+Ppr in the non-heated and heated spent medium of Pfl. Under non-heated treatments, 5 independent cells were measured for Cre monoculture and 10 for Cre+Ppr coculture. Under heated treatment, 12 and 10 independent cells, respectively, were measured. **C-D** The experiment was independently repeated. The results of the second dataset are shown in Fig. S9A. **E** Protegencin levels in the Cre+Ppr coculture increase when proteinase K is added to the Pfl spent medium. Protegencin levels were compared between Cre+Ppr cocultures grown in non-heated Pfl supernatant and in Pfl supernatant treated with proteinase K. In TAP medium, 8 independent cells were measured. In *P. fluorescens* spent medium, 10 independent cells were measured. For the proteinase K-treated *P. fluorescens* spent medium, 9 independent cells were measured. Replicate 2 of the proteinase K treatment (n=2) is shown in Fig. S9B. **C-E** Protegencin levels were quantified by Raman microspectroscopy using the ratio of the triple-bond signal at (2170 ± 100) cm⁻¹ to the reference signal at (1635 ± 80) cm⁻¹. The data are presented as box-and-whisker plots, with the center line indicating the median, the box representing the interquartile range (IQR), and the whiskers extending to the minimum and maximum values. Individual values are shown as scattered points, including both inner and outlier points. Statistical significance was determined using an unpaired two-tailed Welch’s *t*-test (unequal variances). Significance levels are indicated as follows: \*\**P* < 0.01, \*\*\**P* < 0.001 and n.s.: not significant. Exact *P*-values, *t*-values and degrees of freedom are reported in Table S8A, B, D, respectively.

Based on these observations, we discussed different scenarios (Fig. 6B): (i) active secretion of enzyme(s) by *P. fluorescens* or of other heat-sensitive compound(s) that degrade protegencin, and/or (ii) release of intracellular enzyme(s)/compound(s) following lysis of *P. fluorescens* cells caused by *P. protegens*. Although it may be challenging to distinguish between lysis and active secretion, we aimed to know whether bacterial cells and spent medium would contain compound(s)/enzyme(s) that can degrade protegencin.

For this purpose, cocultures of *C. reinhardtii* and *P. protegens* were first incubated with a *P. fluorescens* protein extract, and protegencin levels were subsequently quantified by Raman microspectroscopy. Addition of the *P. fluorescens* protein extract reduced protegencin levels to background levels, comparable to those observed in the pure-cell control (Fig. 6C, S9A; Table S8A, C). Heat treatment of the protein extract resulted in higher protegencin levels than those observed with the untreated extract (Fig. 6C, S9A; Table S8A, C), corroborating that the protective activity is heat sensitive. These data indicate that the protegencin lytic activity is present inside the bacterial cells.

We next tested the effect of *P. fluorescens* spent medium on the protegencin levels. Addition of the spent medium reduced protegencin levels in the *C. reinhardtii–P. protegens* coculture (Fig. 6D, S9A; Table S8B, C). Heat treatment of the spent medium abolished again this effect, resulting in higher protegencin levels (Fig. 6D, S9A; Table S8B, C). The reduction of protegencin in the spent medium was less pronounced than that observed with the cellular protein extract (Fig. 6C-D). Together, these data suggest that the protective activity involves at least one heat-sensitive factor present both extracellularly and within *P. fluorescens* cells. The active factor(s) may be more abundant or concentrated in the cellular fraction than in the extracellular environment, supporting the possibility that they may originate from *P. fluorescens* cells and are being released upon cell lysis. Thus, the active factor(s) would appear automatically in spent medium upon cell lysis of *P. fluorescens,* and a specific secretion process may not be needed.

To examine whether the protective activity is indeed mediated by enzyme(s) or other heat-sensitive compound(s), we treated the *P. fluorescens* spent medium with proteinase K before inoculating the *C. reinhardtii–P. protegens* coculture. Proteinase K treatment resulted in significantly higher protegencin levels than in the untreated spent medium (Fig. 6E, S9B, Table S8D), showing that proteolytic digestion abolishes the protective activity. Together with the heat-treatment experiments, these results strongly support the conclusion that the protective factor produced by *P. fluorescens* is proteinaceous in nature and most likely an enzyme.

Protective interactions against antagonistic microorganisms have been described in several host-associated microbial tripartite systems. However, the underlying mechanisms differ substantially from those identified here. In the *C. reinhardtii–P. protegens* system, it was shown that *Mycetocola lacteus* protects the alga on long term (more than 20 days) by enzymatically cleaving the lipopeptide toxin orfamide A, thereby abolishing its algicidal activity^23^. This interaction is mutualistic, with the bacterium benefiting from algal-derived metabolites while protecting its host. Another study described a distinct mechanism in which a lichen-like association between *C. reinhardtii* and *Aspergillus nidulans* protects the alga from *Streptomyces iranensis* through formation of a stable multicellular consortium^41^. Here, we show a functional shift of a bacterium involved in algal-bacterial interactions. Such a context-dependent functional shift in algal–bacterial interactions was described before for the marine microalga *Emiliania huxleyi* and its associated bacterium *Phaeobacter gallaeciensis*. In this case, the age of the alga plays an important role. During algal ageing, the algal-derived compound *p*-coumaric acid induces a switch in bacterial secondary metabolism from the production of growth-promoting compounds to the algicidal roseobacticide(s), converting the bacterium into an opportunistic pathogen^42^. Our study describes another context-dependent outcome involving three interacting partners. In pairwise cultures, all three partners form amensalistic(-like) relationships, in which neither partner benefits from the interaction: *P. protegens* negatively affects both *C. reinhardtii* and *P. fluorescens*, while *P. fluorescens* negatively affects *C. reinhardtii* (Fig. 7). In the tripartite community, however, the outcome changes markedly. *P. protegens* remains antagonistic towards both *C. reinhardtii* and *P. fluorescens*, whereas *P. fluorescens* inactivates protegencin produced by *P. protegens*, thereby protecting *C. reinhardtii* from its toxicity (Fig. 7).

**Fig. 7.**
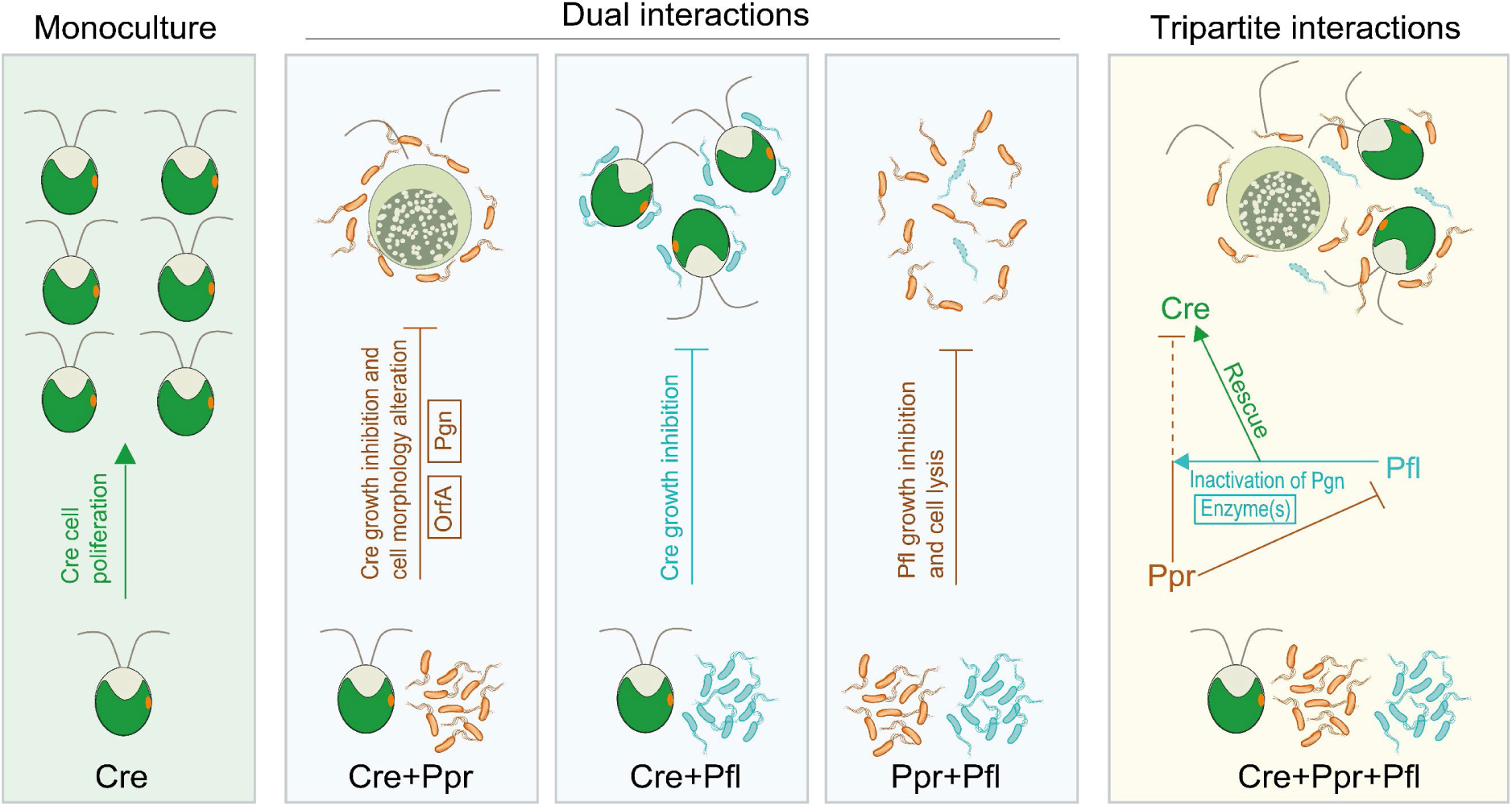
From amensalism to conditional protection. Overview of the bi- and tripartite interactions among *C. reinhardtii* (Cre), *P. protegens* (Ppr), and *P. fluorescens* (Pfl) compared to an algal monoculture. In the bipartite culture of Cre and Ppr, Ppr strongly inhibits algal growth through the production of toxins, including orfamide A and protegencin, resulting in altered algal morphology and cell lysis. However, Ppr does not benefit from Cre, at least in the used TAP medium, resulting in an amensalistic-like relationship between the two microorganisms. In the bipartite culture of Cre and Pfl, Pfl forms an amensalistic interaction with Cre by suppressing algal growth without benefiting from the alga. In the bacterial coculture, Ppr strongly suppresses the growth of Pfl and induces lysis of Pfl cells, while Ppr does not benefit from the presence of Pfl, forming another amensalistic relationship. In the tripartite culture, Ppr remains antagonistic towards Pfl, whereas Pfl produces enzyme(s) that inactivate protegencin, thereby protecting Cre from Ppr-mediated antagonism and enabling algal recovery.

Together, our study expands the current understanding of microbial interactions by emphasizing that the ecological role of a microorganism can depend strongly on its community context: In the presence of the antagonist *P. protegens*, *P. fluorescens*, the enemy of *C. reinhardtii*, is getting transformed into an algal protector. This context-dependent shift highlights the importance of investigating microbial interactions beyond pairwise systems and illustrates how complex communities can generate unexpected protective outcomes through the interplay of amensalism and toxin neutralization.

## Methods

### Strains

Experiments were carried out with *Chlamydomonas reinhardtii* wild-type strain SAG 73.72 (mt+) obtained from the Culture Collection of Algae at Göttingen University (SAG). It is an isolate of Tsubo’s strain C8, which originates from 21 gr (Sager) deriving directly from the originally isolated field strain from Smith, 1945^16^. The bacterium *Pseudomonas protegens* Pf-5 BAA-477™ (named *P. protegens*) was obtained from the American Type Culture Collection (ATCC). *Pseudomonas fluorescens* HKI0770 (named *P. fluorescens*) was provided by the Leibniz Institute for Natural Product Research and Infection Biology, Hans Knöll Institute (HKI), Jena, Germany^28^. For performing FISH, the fluorescently labeled strain *P. fluorescens* HKI0770::MRE-TN7-145 (named *P. fluorescens*\*), also provided by HKI, was used^43^.

### Correlation between OD_600_ and bacterial cell density of *P. protegens* and *P. fluorescens*

The correlation curve between OD_600_ and bacterial cell density for *P. protegens* was adopted from Carrasco Flores *et al*. (2024)^23^. To establish a calibration curve for *P. fluorescens*, colony-forming units (CFUs) were counted and correlated with the corresponding OD_600_ values (Fig. S2A).

### Algal and bacterial mono- and cocultures

The mono- and coculture experiments were carried out according to Carrasco Flores *et al*. (2024)^23^. *C. reinhardtii* pre-, mono- and cocultures were grown in liquid Tris-Acetate-Phosphate (TAP) medium^16^ at 23 °C under a 12:12 light-dark cycle under white light (Osram L36W/840, lumilux, cool white, OSRAM, Germany) with a light intensity of about 60 μmol m^−2^ s^−1^ and orbital shaking (80 rpm). Bacterial precultures were grown overnight at 26 °C with shaking at 80 rpm in Luria-Bertani (LB) broth (Carl Roth, Germany) for *P. protegens* and Standard Medium (SM/5; Formedium, England) for *P. fluorescens*. Before cocultivation, bacterial cells were washed twice with TAP medium and mixed with *C. reinhardtii*. Bipartite algal-bacterial cultures were initiated at a density of 1×10⁵ algal cells ml⁻¹ and 2.5×10⁷ bacterial cells ml⁻¹, corresponding to an algae-to-bacteria ratio of 1:250^23^. For bacterial bipartite cultures, *P. protegens* and *P. fluorescens* were each inoculated in TAP at 2.5×10⁷ cells ml⁻¹, resulting in a 1:1 bacterial ratio. Tripartite cultures were established in TAP with final densities of 1×10⁵ *C. reinhardtii* cells ml⁻¹ and 2.5×10⁷ cells ml⁻¹ of each bacterial species, corresponding to a final ratio of 1:250:250 (*C. reinhardtii*: *P. protegens*: *P. fluorescens*). Cocultures were incubated as described above for the monoculture of *C. reinhardtii* over a period of 10 days. Photographic documentation of cultures was performed daily. All coculture experiments were independently repeated three times. Along with each biological replicate, three technical replicates were performed, unless otherwise specified.

### Fluorescence *in situ* hybridization (FISH)

Tripartite cultures were established as described above using the fluorescently labeled *P. fluorescens*\* strain described in the Strain section. The cells were harvested and washed twice with phosphate-buffered saline (PBS; 137 mM NaCl, 2.7 mM KCl, 10 mM Na_2_HPO_4_, 1.8 mM KH_2_PO_4_, pH 7.4) and fixed using 4% paraformaldehyde for 4 h at 4 °C. FISH was performed using the probes, EUK516 (for eukaryotes; probeBase accession no. pB-162) labeled with Alexa Fluor™ 488 dye (ThermoFisher Scientific, USA), GAM42a (for Gammaproteobacteria; probeBase accession no. pB-174) labeled with Alexa Fluor™ 647 dye (ThermoFisher Scientific, USA) and PSE227 (*Pseudomonas*-specific; probeBase accession no. pB-2539) labeled with ATTO 465 dye (Eurofins Genomics, Germany).

The FISH protocol was performed as previously described by Suma and Stallforth (2025)^43^, with modifications based on Zwirglmaier (2010)^44^. A 10 μl of each culture was spread on a microscope slide with wells (Paul Marienfeld, Germany). After air drying, sequential ethanol dehydration (50%, 80%, and 100%) was performed. The slide was then incubated with hybridization buffer (1 pmol ml^−1^ EUK516, GAM42a and PSE227 probes, 20 mM Tris–HCl, 0.9 M NaCl, 0.01% SDS, and 35% formamide, pH 7.5) at 46 °C for 2.5 h. After hybridization, the slide was placed in pre-warmed wash buffer (20 mM Tris–HCl, 0.7 M NaCl, 5 mM Na_2_EDTA) for 20 min at 48 °C. All chemicals were purchased from Carl Roth, Germany. The slide was then rinsed with ice-cold water. After drying, the cells were mounted in ProLong™ Glass antifade reagent (ThermoFisher Scientific, USA) and a coverslip was placed over the slide.

Imaging was performed using a spinning disk confocal laser scanning microscope (AxioObserver.Z1/7, Carl Zeiss, Germany) equipped with a 100×/1.40 NA Plan-Apochromat oil-immersion DIC M27 objective lens (Carl Zeiss, Germany). Images were captured using diode lasers: 445 nm (40 mW), 488 nm (50 mW), 561 nm (50 mW), and 638 nm (75 mW) with the corresponding emission bandpass (BP) filters: BP 450/50, 485/30, BP 600/50, and BP 629/62. Images were processed using ZEN 2.6 (Blue edition, Carl Zeiss, Germany) imaging software.

### Acetic acid assay^45^

Acetate concentrations in mono-, bi-, and tripartite cultures of *C. reinhardtii*, *P. protegens*, and *P. fluorescens* were measured daily over five days using an acetic acid assay kit (Megazyme, Ireland) according to the manufacturer’s instructions.

### Acetic acid-dependent and -independent tripartite cultures

Tripartite cultures were established as described above. On day 2 of incubation, after acetate depletion, acetate was replenished to a final concentration of 1.04 g l⁻¹ in one set of cultures, whereas a second set was maintained without acetate supplementation. To maintain a constant pH, acetate was added as Tris-acetate buffer (pH 7) at the same concentration as in fresh TAP medium^16^.

### Cell quantification of *P. protegens* and *P. fluorescens* via plating

Quantification of the cell numbers of *P. protegens* and *P. fluorescens* in mono- and cocultures with *C. reinhardtii*, or in tripartite cultures was performed by plating, as the two bacterial species exhibit distinct colony morphologies on SM/5 agar plates (Fig. S2B). Serial dilutions were prepared in dH_2_O, and each sample was plated in three technical replicates. Plates were incubated at 28 °C for 48 hours until colonies were clearly visible. The experiment was conducted daily over 10 days to generate growth curves. Bacterial cell density was determined by colony-forming unit (CFU).

### *C. reinhardtii* cell quantification via counting

Algal cell density was determined daily using an improved Neubauer cell counting chamber^23^.

### Microscopy of bacteria

Lattice-SIM images were acquired on a Zeiss Elyra 7 microscope using a Plan-Apochromat 63×/1.4 Oil DIC M27 objective (Carl Zeiss Microscopy GmbH, Germany). Illumination was provided by the Elyra’s 405 nm laser line (Carl Zeiss Microscopy GmbH, Germany). The 405 nm laser line was used to excite the autofluorescence of *P. fluorescens*, whereas *P. protegens* exhibited only very low residual levels of autofluorescence under these conditions. For each individual image, 13 phase images were recorded. SIM reconstruction was performed in ZEN Blue 3.11 with the “baseline cut” option deactivated. The final pixel size in reconstructed SIM images is 30 nm. Transmission images of bacteria have been recorded using the same objective and the Elyra 7 in widefield transmission mode with illumination provided by a HAL100 halogen lamp (Carl Zeiss Microscopy GmbH, Germany).

### Quantification of bacterial cell debris

For statistical evaluation of bacterial cell debris, a 5 × 5 tile scan (10% overlap) resulting in a final image size of 380.20 µm ×379.75 µm was acquired to cover a large area of the samples. Cellular debris was quantified in the background of images from the *P. protegens*–*P. fluorescens* coculture, with monocultures of each bacterium serving as controls. For each sample, debris was quantified across all 25 tiles of the scan. The resulting values were normalized to the debris abundance in the *P. fluorescens* monoculture, which was set to 1.

### *C. reinhardtii* and *P. protegens* cocultures grown in spent medium of a tripartite culture

100 ml of a tripartite culture of *C. reinhardtii*, *P. protegens* and *P. fluorescens* was prepared as described above. On day 3, the culture supernatant was collected after centrifugation at 4000 × *g* for 30 min. To eliminate remaining live bacterial cells, the supernatant was exposed to UV-C light for 1 h using a Philips TUV 30W/G30T8 lamp (253.7 nm; UV-C output, 12.3 W; Signify, Netherlands). The supernatant, indicated as spent medium, was then divided into two equal portions. One portion was heat-treated at 121 °C for 35 min by autoclaving, while the other remained untreated. As nutrients in the spent medium had already been depleted, all TAP medium components^16^ were replenished to their original concentrations before inoculation. Both non-heated and heated spent media were subsequently used to cultivate *C. reinhardtii* monocultures and cocultures of *C. reinhardtii* and *P. protegens*. The cocultures in fresh TAP were used as controls.

### Raman sample preparation^46^

Raman samples were prepared according to Vuong *et al*. (2025)^46^. Briefly, cells were fixed in 4% (v/v) formalin, embedded in 1% (w/v) agarose on CaF_2_ substrates, and stored at 4 °C until Raman microspectroscopy.

### Raman measurements

Raman microspectroscopy of algal samples was performed following Vuong *et al.* (2025)^46^ using the identical 785 nm Raman microscopy setup (WITec alpha300, WITec Wissenschaftliche Instrumente und Technologie GmbH, Germany) and water-dipping objective (60×/1.0 NA NIR, Nikon, Japan). Briefly, individual cells were imaged using an excitation power of 100 mW before the objective (∼80 mW at the objective’s front lens) and a spatial resolution of 3 px µm⁻¹, representing slight oversampling relative to Abbe’s limit. Following Vuong *et al.* (2025)^46^, each position was measured twice, with an initial 0.1 s px⁻¹ bleaching scan, followed by a 0.5 s px⁻¹ acquisition scan for Raman signal recording. Only the latter measurement was used for further analysis. Image sizes were adjusted to encompass the entire cell (cluster) with a small safety margin, resulting in sampled areas of approximately (10 µm)²–(20 µm)².

### Raman data analysis

The generated datasets were processed in *Python* 3.13.5 using the scripts provided via https://doi.org/10.5281/zenodo.22737390. Pre-processing comprised spike removal, piecewise SNIP^47,48^ background correction, and wavenumber correction. Algal cell and background regions were separated by *k*-means clustering, followed by, if necessary, manual adjustments in *ImageJ*. The corrected spectra were integrated pixelwise at (2170±100) cm⁻¹ after subtraction of an additional linear baseline and at (1635±80) cm⁻¹ and (2920±150) cm⁻¹ relative to the zero-intensity baseline. Individual algal cells were delineated in clusters by watershed segmentation informed by the spatial intensity distribution of the combined 1635 and 2920 cm⁻¹ Raman maps. Parameters were adjusted to best match the majority of measured cell positions with clusters. Where over-segmentation occurred, the corresponding regions were manually merged in *ImageJ*.

The protegencin content was estimated as stated in the axes labels of the figures by dividing the triple bond stretching band at (2170 ± 100)_bg_ cm⁻¹ against the amide 1 vibration at (1635±80) cm⁻¹.

### Preparation of *P. fluorescens* spent medium and protein extract^23^, including heat- and proteinase K treatment

An overnight *P. fluorescens* preculture grown in SM/5 medium at 28 °C was subcultured into fresh medium for 2 days, washed in TAP twice and transferred to TAP medium for another 2 days, and centrifuged (4000 × *g*, 30 min, 4 °C). 60 ml of the resulting supernatant was collected and supplemented with TAP nutrients to restore their original concentrations before inoculation^16^. The cell pellet was used for protein extraction on ice as described by Carrasco Flores *et al*. (2024)^23^. After extraction, fresh TAP medium was added to the protein extract to a final volume of 60 ml, matching the volume of the supernatant used for the coculture experiments.

The supernatant or protein extract was divided into two equal portions. One portion was heat-treated by autoclaving at 121 °C for 35 min, while the other remained untreated. Both treated and untreated samples were used for coculture experiments consisting of a 15 ml algal culture and a 15 ml *C. reinhardtii*-*P. protegens* coculture. In case of proteinase K (P6556, Sigma-Aldrich, Germany) treatment, cocultures of *C. reinhardtii* and *P. protegens* were established using *P. fluorescens* spent medium that had been treated with proteinase K at a final concentration of 50 µg ml⁻¹. As additional controls, identical cocultures were performed in TAP medium alone and in TAP supplemented with proteinase K at the same concentration.

All samples were collected on day 2 and subsequently prepared for Raman spectroscopy analysis as mentioned above.

### Statistical analysis

Statistical analyses were performed using unpaired, two-tailed Welch’s *t*-tests. Exact *P*-values, *t*-values, and degrees of freedom are reported individually in the supplementary tables for each comparison. The n values represent the number of independent measurements or biological replicates are specified in the corresponding figure legends. Data are presented as mean ± SD or as box-and-whisker plots in the case of protegencin data, with the center line indicating the median, the box representing the interquartile range, and the whiskers extending to the minimum and maximum values, as indicated in the figure legends.

## Acknowledgements

We thank Dr. Prateek Shetty for helpful advice and proofreading of the manuscript.

## Author contributions

TV, CS, MS, JP, PS and MM designed the project. TV, CS, HRS, PT and SJ performed experiments, and all authors contributed to the analysis and interpretation of results. TV and MM wrote the paper with input from all coauthors.

## Funding

TV had a fellowship from the Carl Zeiss Foundation under the umbrella of JSMC. PT, JP, PS and MM received funding from the DFG within the Germany’s Excellence Strategy – EXC 2051 – Project ID 390713860. We thank the Microverse Imaging Center also financed by EXC 2051 for providing microscope facility support for data acquisition. The Free State of Thuringia funded the ELYRA 7 with grant no. 2019 FGI 0003. We thank the Werner Siemens-Stiftung (Paleobiotechnology to PS) for funding. SJ and JP received funding from The EuropeanUnionTranscan-3 joint transnational call programme awarded to “ReachGLIO” project (TRANSCAN2022-784-017) under the grant agreement No.964264 and funded by the “The Federal Ministry of Research, Technology and Space (BMFTR)” in Germany (Funding code: 01KT2327).

## Competing interests

The authors declare no competing interests.

## Supplementary materials

**Supplementary Fig. S1 to S9**

**Supplementary Tables S1 to S8**

**Correspondence** and requests for materials should be addressed to Maria Mittag.

## Data availability

A Zenodo upload of the Raman data will be set to public once the paper is published.

## Notes

### Competing Interest Statement

The authors have declared no competing interest.

## References

1. Faust, K. & Raes, J. Microbial interactions: From networks to models. Nature Reviews Microbiology 10, 538–550 (2012).

2. Ponomarova, O. & Patil, K. R. Metabolic interactions in microbial communities: Untangling the Gordian knot. Current Opinion in Microbiology 27, 37–44 (2015).

3. Levine, J. M., Bascompte, J., Adler, P. B. & Allesina, S. Beyond pairwise mechanisms of species coexistence in complex communities. Nature 546, 56–64 (2017).

4. Maberly, S. C. & Gontero, B. Ecological imperatives for aquatic CO_2_-concentrating mechanisms. Journal of Experimental Botany 68, 3797–3814 (2017).

5. Gorman, L., Harrison, E., Highfield, A., Helliwell, K. & Wheeler, G. Get the message: Uncovering cellular signalling processes in marine phytoplankton. Advances in Marine Biology 104, 85–115 (2026).

6. Field, C. B., Behrenfeld, M. J., Randerson, J. T. & Falkowski, P. Primary production of the biosphere: Integrating terrestrial and oceanic components. Science 281, 237–240 (1998).

7. Kouzuma, A. & Watanabe, K. Exploring the potential of algae/bacteria interactions. Current Opinion in Biotechnology 33, 125–129 (2015).

8. Cooper, M. B. & Smith, A. G. Exploring mutualistic interactions between microalgae and bacteria in the omics age. Curr. Opin. Plant Biol. 26, 147–153 (2015).

9. Burgunter-Delamare, B., Shetty, P., Vuong, T. & Mittag, M. Exchange or eliminate: the secrets of algal-bacterial relationships. Plants 13, 829 (2024).

10. Torres, M. J. et al. Chlamydomonas-*Methylobacterium oryzae* cooperation leads to increased biomass, nitrogen removal and hydrogen production. Bioresour. Technol. 352, 127088 (2022).

11. Hom, E. F. Y., Aiyar, P., Schaeme, D., Mittag, M. & Sasso, S. A chemical perspective on microalgal-microbial interactions. Trends Plant Sci. 20, 689–693 (2015).

12. Seymour, J. R., Amin, S. A., Raina, J. B. & Stocker, R. Zooming in on the phycosphere: The ecological interface for phytoplankton-bacteria relationships. Nature Microbiology 2, 17065 (2017).

13. Helliwell, K. E., Shibl, A. A. & Amin, S. A. The diatom microbiome: New perspectives for diatom-bacteria symbioses. The Molecular Life of Diatoms. 679–712 (Springer International Publishing, 2022).

14. Brodie, J. et al. Biotic interactions as drivers of algal origin and evolution. New Phytologist 216, 670–681 (2017).

15. Calatrava, V., et al. *Chlamydomonas reinhardtii*, a reference organism to study algal– microbial interactions: why can’t they be friends? Plants 12, 788 (2023).

16. Harris. The Chlamydomonas Sourcebook: A Comprehensive Guide to Biology and Laboratory Use. (Academic Press Inc, 1989).

17. Lingappa, U. F. et al. Field strains of the unicellular alga *Chlamydomonas reinhardtii* exhibit multicellular characteristics that shape their interactions. ISME Journal 20, 1–15 (2026).

18. Barrett, M. R. & Koch, A. R. Effects of ammonium sulfate and urea on the growth of *Chlorophycean* algae from rice fields. J. Environ. Qual. 11, 187–191 (1982).

19. Ghasemi, Y. et al. Characterization of hydrocortisone biometabolites and 18S rRNA gene in *Chlamydomonas reinhardtii* cultures. Molecules 13, 2416–2425 (2008).

20. Lin, C. S., Chou, T. L. & Wu, J. T. Biodiversity of soil algae in the farmlands of mid-Taiwan. Bot. Stud. 54, 1–12 (2013).

21. Durán, P. et al. Shared features and reciprocal complementation of the *Chlamydomonas* and *Arabidopsis* microbiota. Nat. Commun. 13, 406 (2022).

22. Aiyar, P. et al. Antagonistic bacteria disrupt calcium homeostasis and immobilize algal cells. Nat. Commun. 8, 1756 (2017).

23. Carrasco Flores, D., et al. A mutualistic bacterium rescues a green alga from an antagonist. Proc. Natl. Acad. Sci. U. S. A. 121, e2401632121 (2024).

24. Jiamin, S., Xinxin, Z., Yucong, W., Guoqing, C. & Guozhong, F. Efficient production of pyrrolnitrin by optimizing culture medium and blocking competitive secondary metabolic pathways in *Pseudomonas protegens* JP2-4390. Rice Science 32, 156–159 (2025).

25. Hotter, V. et al. A polyyne toxin produced by an antagonistic bacterium blinds and lyses a Chlamydomonad alga. Proc. Natl. Acad. Sci. U. S. A. 118, e2107695118 (2021).

26. Hou, Y. et al. A cyclic lipopeptide produced by an antagonistic bacterium relies on its tail and transient receptor potential-type Ca^2+^ channels to immobilize a green alga. New Phytologist 237, 1620–1635 (2023).

27. Rose, M. M. et al. The bacterium Pseudomonas protegens antagonizes the microalga *Chlamydomonas reinhardtii* using a blend of toxins. Environ. Microbiol. 23, 5525–5540 (2021).

28. Klapper, M., Götze, S., Barnett, R., Willing, K. & Stallforth, P. Bacterial alkaloids prevent amoebal predation. Angew. Chem. Int. Ed. 55, 8944–8947 (2016).

29. Suman, B., Vijaya Gopal, A., Subhash Reddy, R. & Triveni, S. Isolation and characterization of *Pseudomonas fluorescens* from rice rhizospheric soils of Rangareddy district in Telangana state. J. Pure Appl. Microbiol. 9, 339–344 (2015).

30. Sharifi Noori, M. S. & Mohd Saud, H. Potential plant growth-promoting activity of *Pseudomonas* sp isolated from paddy soil in Malaysia as biocontrol agent. J. Plant Pathol. Microbiol. 03, 1–4 (2012).

31. Hori, T., Noll, M., Igarashi, Y., Friedrich, M. W. & Conrad, R. Identification of acetate-assimilating microorganisms under methanogenic conditions in anoxic rice field soil by comparative stable isotope probing of RNA. Appl. Environ. Microbiol. 73, 101–109 (2007).

32. Shen, L. D. et al. Evidence for the cooccurrence of nitrite-dependent anaerobic ammonium and methane oxidation processes in a flooded paddy field. Appl. Environ. Microbiol. 80, 7611–7619 (2014).

33. Roca, C. & Olsson, L. Dynamic responses of *Pseudomonas fluorescens* DF57 to nitrogen or carbon source addition. J. Biotechnol. 86, 39–50 (2001).

34. Perrineau, M. M. et al. Using natural selection to explore the adaptive potential of *Chlamydomonas reinhardtii*. PLoS One 9, e92533 (2014).

35. Balthazar, C. et al. Pyoluteorin and 2,4-diacetylphloroglucinol are major contributors to *Pseudomonas protegens* Pf-5 biocontrol against Botrytis cinerea in cannabis. Front. Microbiol. 13, 945498 (2022).

36. Gu, Q., et al. The role of pyoluteorin from Pseudomonas protegens Pf-5 in suppressing the growth and pathogenicity of Pantoea ananatis on maize. Int. J. Mol. Sci. 23, 6431 (2022).

37. Quecine, M. C. et al. An interspecies signaling system mediated by fusaric acid has parallel effects on antifungal metabolite production by *Pseudomonas protegens* strain Pf-5 and antibiosis of *Fusarium* spp. Appl. Environ. Microbiol. 82, 1372–1382 (2016).

38. Mwanza, C. et al. Polyyne production is regulated by the transcriptional regulators PgnC and GacA in *Pseudomonas protegens* Pf-5. Appl. Environ. Microbiol. 91, 22 (2025).

39. Laakso, S., Soderling, E. & Nurmikko, V. Methionine degradation by *Pseudomonas fluorescens* UK1 and its methionine utilizing mutant. J. Gen. Microbiol. 94, 305–312 (1976).

40. Liao, R. Z., Yu, J. G. & Himo, F. Mechanism of tungsten-dependent acetylene hydratase from quantum chemical calculations. Proc. Natl. Acad. Sci. U. S. A. 107, 22523–22527 (2010).

41. Krespach, M. K. C. et al. Lichen-like association of *Chlamydomonas reinhardtii* and *Aspergillus nidulans* protects algal cells from bacteria. ISME Journal 14, 2794–2805 (2020).

42. Seyedsayamdost, M. R., Case, R. J., Kolter, R. & Clardy, J. The Jekyll-and-Hyde chemistry of *Phaeobacter gallaeciensis*. Nat. Chem. 3, 331–335 (2011).

43. Suma, H. R. & Stallforth, P. Pleiotropic regulation of bacterial toxin production and Allee effect govern microbial predator-prey interactions. ISME Communications 5, ycaf031 (2025).

44. Zwirglmaier, K. Detection of prokaryotic cells with fluorescence in situ hybridization. Methods Mol. Biol. 659, 349–362 (2010).

45. Shetty, P. et al. Multiomics studies reveal how ambient temperature changes govern cellular responses of *Chlamydomonas*. Plant Cell 38, 1–23 (2026).

46. Vuong, T. et al. Metamorphosis of a unicellular green alga in the presence of acetate and a spatially structured three-dimensional environment. New Phytologist 245, 1180– 1196 (2025).

47. Ryan, C. G., Clayton, E., Griffin, W. L., Sie, S. H. & Cousens, D. R. SNIP, a statistics-sensitive background treatment for the quantitative analysis of PIXE spectra in geoscience applications. *Nuclear Inst. and Methods in Physics Research*, B 34, 396– 402 (1988).

48. Morháč, M., Kliman, J., Matoušek, V., Veselský, M. & Turzo, I. Background elimination methods for multidimensional coincidence γ-ray spectra. Nucl. Instrum. Methods Phys. Res. A 401, 113–132 (1997).

