## Supplemental Figures S1-S9 for "The enemy of my enemy: A bacterial antagonist rescues *Chlamydomonas* by inactivating a toxin from a rival"

The following Supplementary Figures are available for this article:

Supplementary Figures S1-S9

### Supplementary Figures

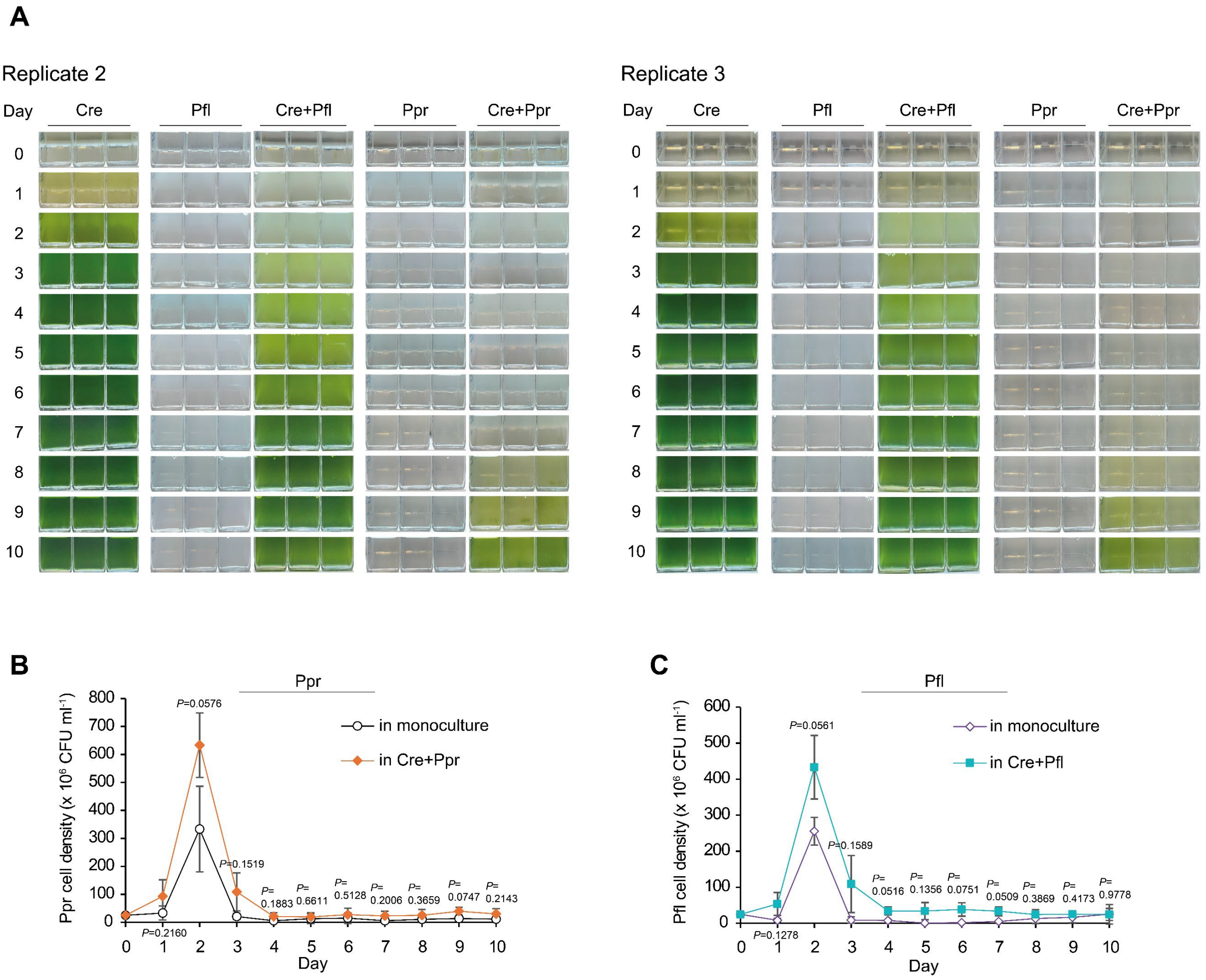


**Fig. S1 | Bipartite cultures of *Chlamydomonas reinhardtii* (Cre) with *Pseudomonas protegens* (Ppr) and *Pseudomonas fluorescens* (Pfl) and bacterial growth. A** Photographs of algal and bacterial monocultures and cocultures. Independent biological replicates 2 and 3 of the bipartite cultures of Cre with Ppr and Pfl are shown. Independent biological replicate 1 is shown in Fig. 1A. **B** The growth of Ppr in the bipartite culture with Cre compared with its monoculture over 10 days. **C** The growth of Pfl in the bipartite culture with Cre compared with its monoculture over 10 days. **B, C** Bacterial cell numbers were determined by colony-forming unit (CFU) assays (see Methods). Values represent the mean of three independent biological replicates (n=3). Error bars indicate the standard deviation (SD). Statistical significance was determined by comparing cell numbers in the bipartite culture with those in the respective monoculture using an unpaired two-tailed Welch's *t*-test (unequal variances). Unlabeled comparisons are not statistically significant. Exact *P*-values, *t*-values and degrees of freedom are reported in Table S2.

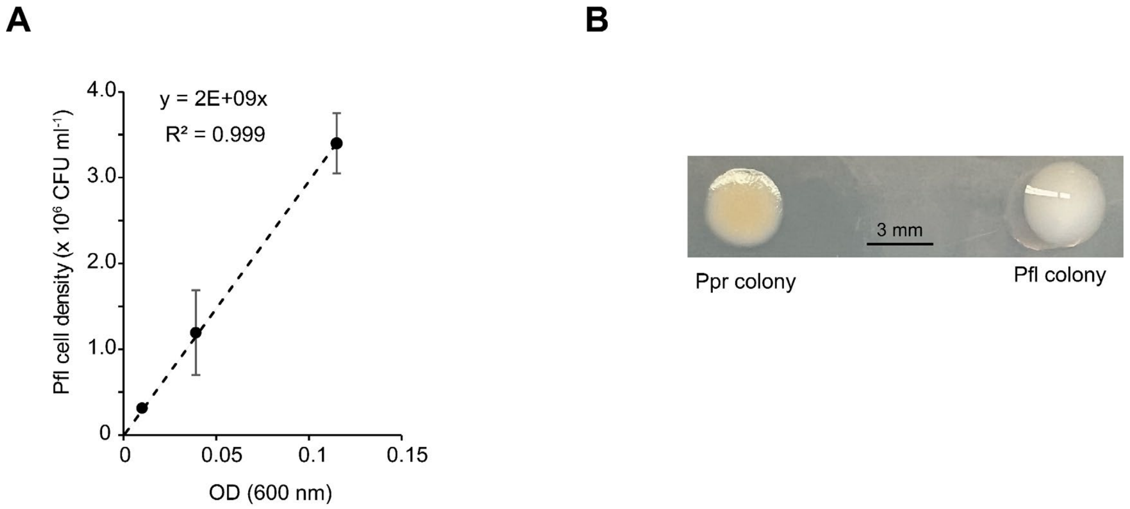


**Fig. S2 | Calibration of bacterial cell density based on $OD_{600}$ (A) and single colonies of *P. protegens* and *P. fluorescens* on SM/5 agar plate (B)**. **A** Correlation between $OD_{600}$ values and *P. fluorescens* (Pfl) cell density. An overnight Pfl culture was grown in SM/5 medium at 26 °C with shaking. $OD_{600}$ was measured for a series of diluted bacterial suspensions using a Jenway 6305 spectrophotometer. For each dilution, 100 µl was plated on SM/5 agar in three technical replicates and incubated at 26 °C for 2 days. Colony-forming units (CFUs) were counted and used to generate the $OD_{600}$-cell density calibration curve. The calibration curve for *P. protegens* was used from Carrasco Flores *et al*. (2024). **B** Representative single colonies of *P. protegens* (Ppr) and Pfl on SM/5 agar plates.

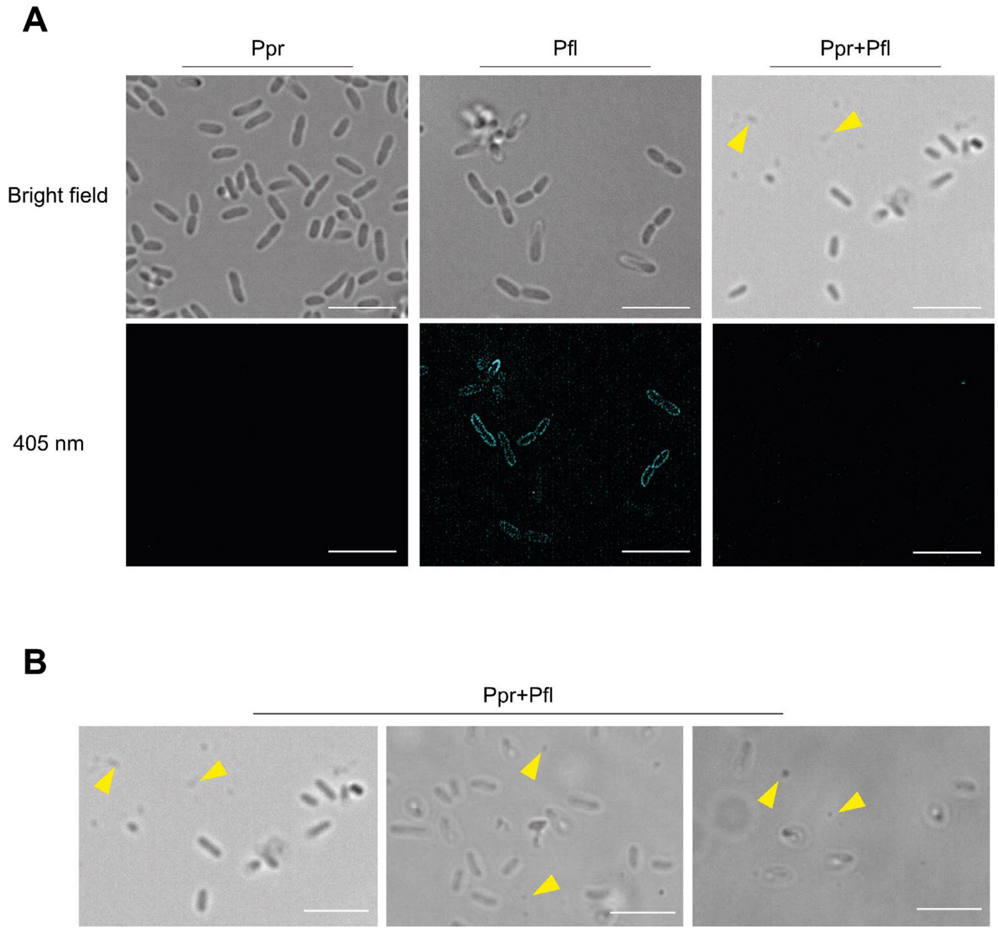


**Fig. S3 | *P. protegens* (Ppr) induces lysis of *P. fluorescens* (Pfl) in coculture. A** Representative images of Ppr and Pfl in their mono- and cocultures. Pfl was identified by its autofluorescence upon 405 nm excitation (turquoise channel), whereas Ppr showed only very low residual fluorescence under these conditions. **B** Additional representative brightfield images of debris observed in Ppr+Pfl cocultures**. A-B** Images were acquired using a Zeiss Elyra microscope (see Methods). Yellow arrows indicate Pfl cell debris. Scale bars = 5 µm.

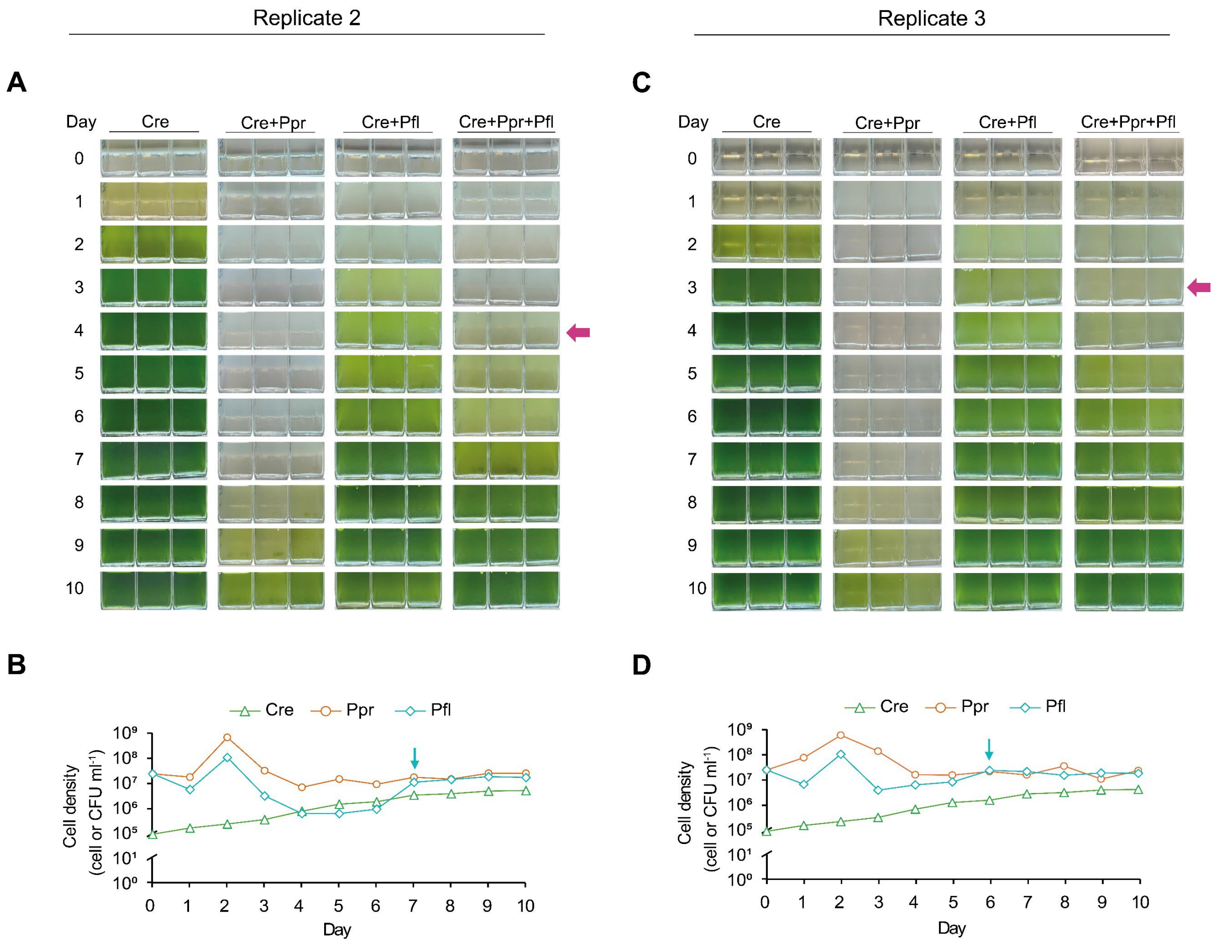


**Fig. S4 | Tripartite culture of *C. reinhardtii* (Cre) with *P. protegens* (Ppr) and *P. fluorescens* (Pfl) reveal a protectory role of Pfl. A, C** Tripartite cultures of Cre, Ppr and Pfl photographed over 10 days. The magenta arrow indicates recovery of Cre. **B, D** Growth curves of Cre, Ppr and Pfl in their tripartite culture over 10 days. The turquoise arrow indicates the recovery of Pfl. **A-B** represents independent biological replicate 2; **C-D** represents biological replicate 3. Independent biological replicate 1 is shown in Fig. 3A, B.

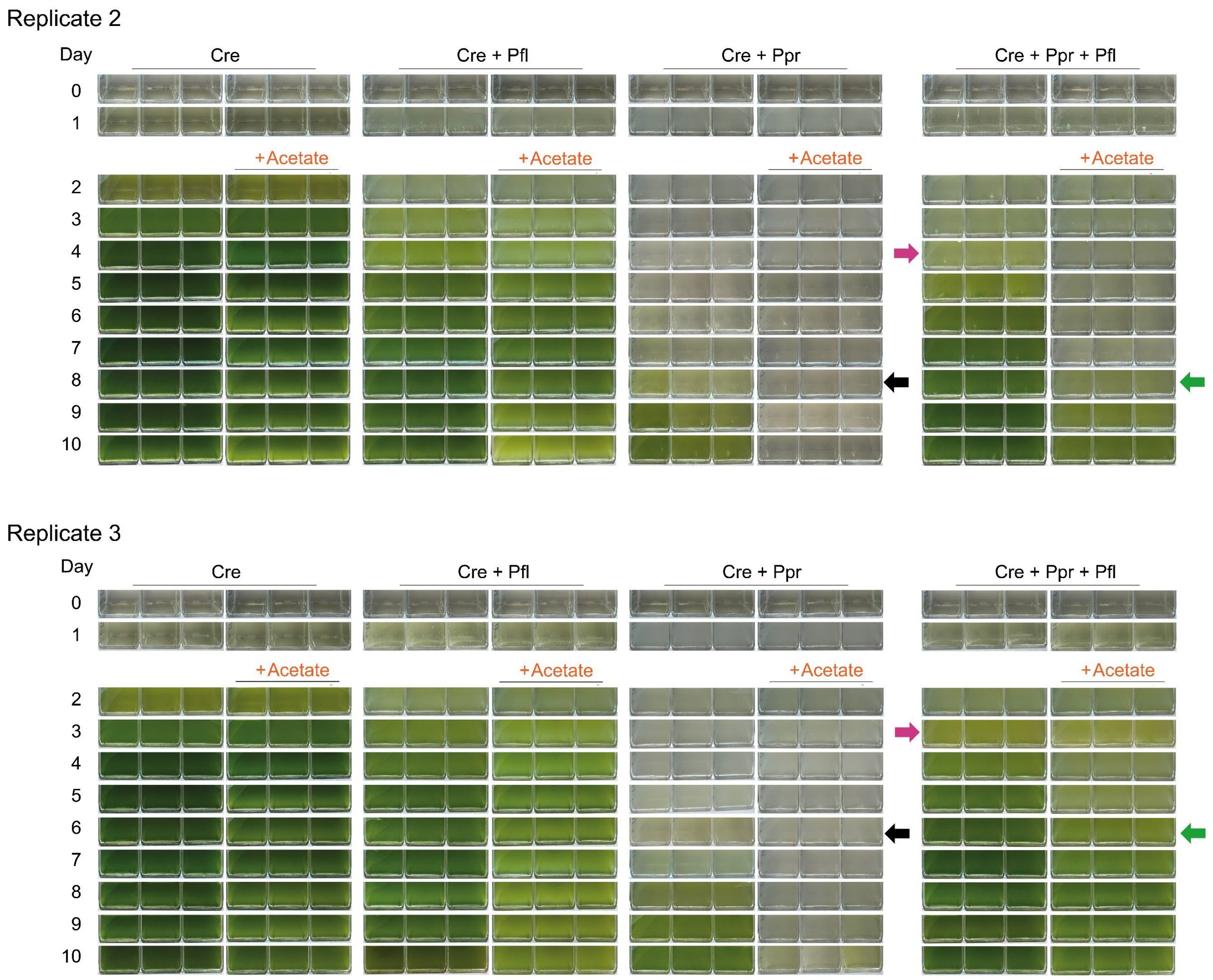


**Fig. S5 | Acetate independence assay in the tripartite culture of *C. reinhardtii* (Cre), *P. protegens* (Ppr), and *P. fluorescens* (Pfl).** Independent biological replicates 2 and 3 are shown. Independent biological replicate 1 is present in Fig. 4B. Cre monocultures and Cre+Ppr and Cre+Pfl cocultures served as controls. Six flasks were prepared for each condition, with acetic acid, buffered at pH 7, added (+acetate) to three flasks on day 2; the remaining flasks received no additional acetate. Magenta and green arrows indicate recovery of Cre, while the black arrow marks consistent bleaching of the Cre+Ppr coculture.

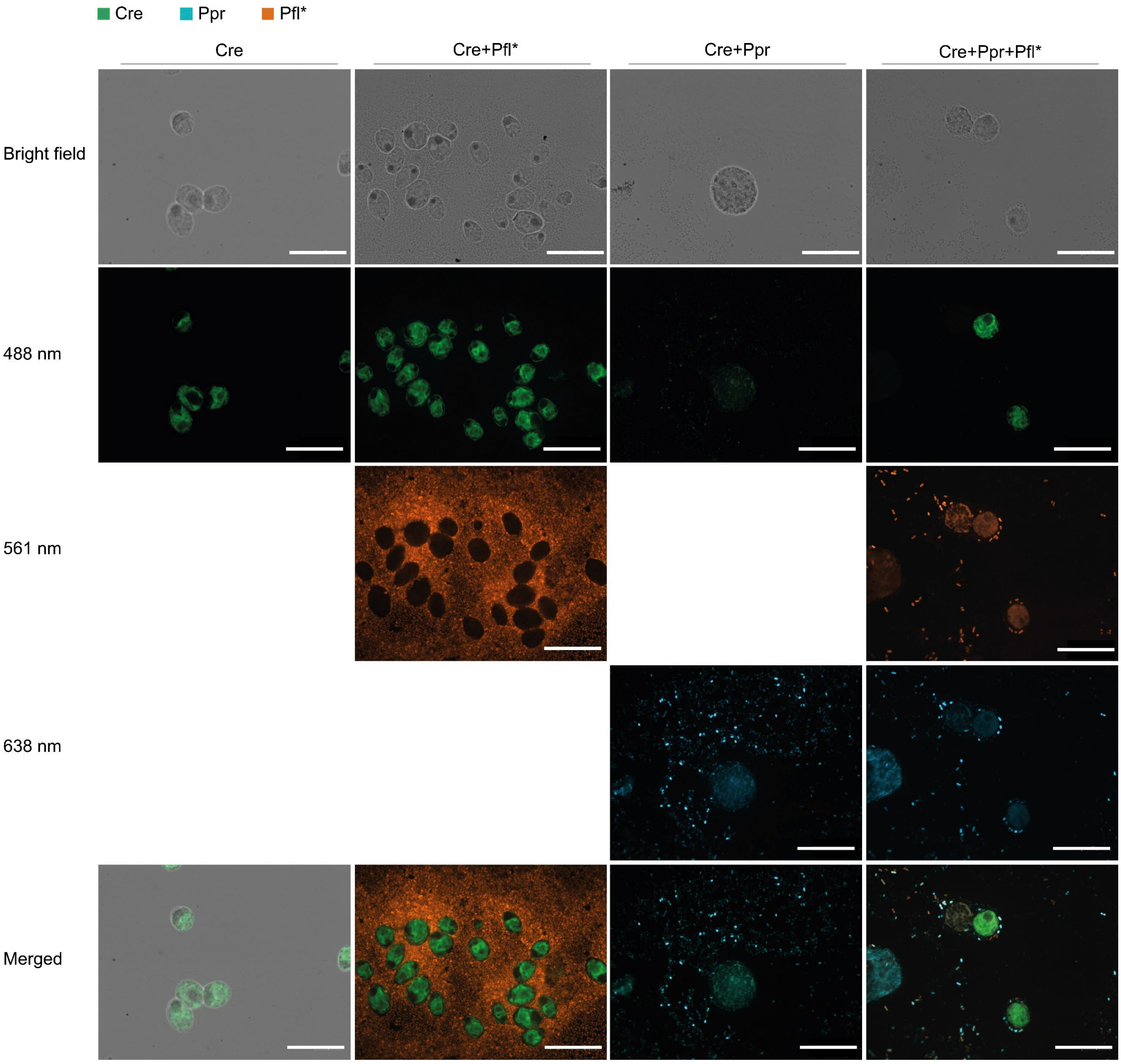


**Fig. S6 | Overview of FISH images of different cocultures of *C. reinhardtii* (Cre) with *P. protegens* (Ppr) and *P. fluorescens** (Pfl*).** The strain *P. fluorescens* HKI0770::MRE-TN7-145 (Pfl*) (see Methods) was used in this experiment. Samples include the Cre monoculture, Cre+Pfl* coculture, Cre+Ppr coculture, and Cre+Ppr+Pfl* tripartite culture. Images were acquired using a spinning disk confocal laser scanning microscope (AxioObserver.Z1/7, Carl Zeiss) equipped with a 100x/1.40 NA Plan-Apochromat oil-immersion DIC M27 objective lens (Carl Zeiss). Images were captured using diode lasers: 488 nm (50 mW), 561 nm (50 mW), and 638 nm (75 mW) with the corresponding emission bandpass (BP) filters: BP 450/50, 485/30, BP 600/50, and BP 629/62. Merged images were generated using the 488 nm channel for Cre, the 561 nm channel for Pfl*, and the 638 nm channel for Ppr. Scale bars = 20 µm.

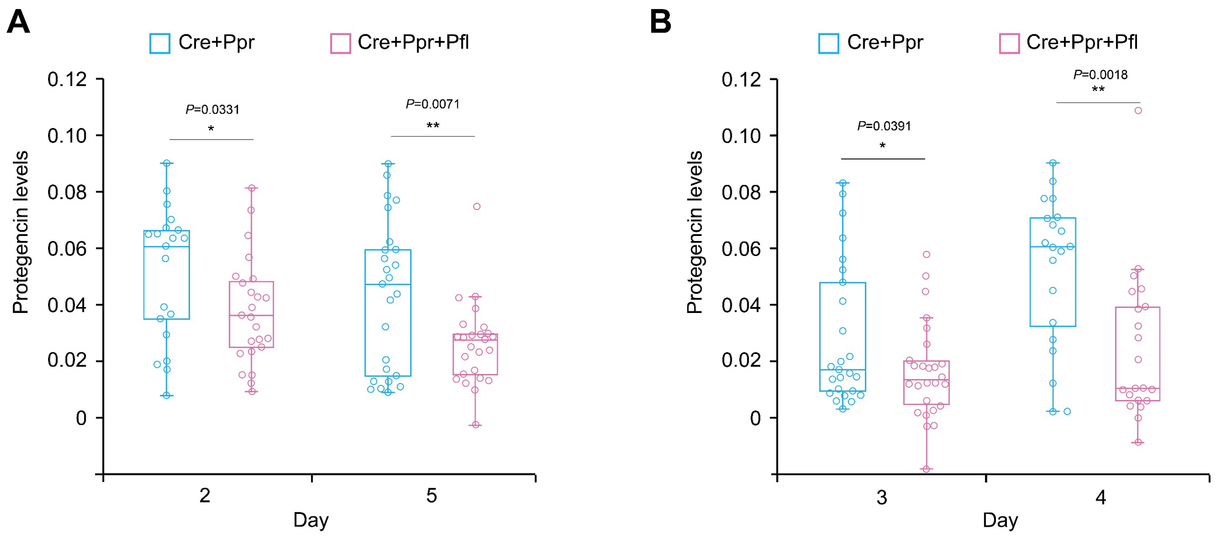


**Fig. S7 | Protegencin levels in the tripartite culture of *C. reinhardtii* (Cre) with *P. protegens* (Ppr) and *P. fluorescens* (Pfl) compared to the coculture of *C. reinhardtii* with *P. protegens* at days 2 and 5 (A) as well as days 3 and 4 (B). A** Data represent biological independent replicate 2. Replicate 1 is shown in Fig. 5B. On day 2, 21 independent cells were measured for the Cre+Ppr bipartite culture and 24 for the Cre+Ppr+Pfl tripartite culture. On day 5, 25 independent cells were measured, respectively, for each condition. **B** On day 3, 25 independent cells were measured for the Cre+Ppr bipartite culture and 26 for the Cre+Ppr+Pfl tripartite culture. On day 4, 20 and 21 independent cells were measured, respectively. **A-B** Protegencin levels were quantified by Raman microspectroscopy using the ratio of the triple-bond signal at (2170 ± 100) $cm^{-1}$ to the reference signal at (1635 ± 80) $cm^{-1}$. For each sample, at least 20 cells were analyzed. The data are presented as box-and-whisker plots, with the center line indicating the median, the box representing the interquartile range (IQR), and the whiskers extending to the minimum and maximum values. Individual values are shown as scattered points, including both inner and outlier points. Statistical significance was determined using an unpaired two-tailed Welch's *t*-test (unequal variances). Significance levels are indicated as follows: $^{**}P < 0.01$, and n.s.: not significant. Exact *P*-values, *t*-values and degrees of freedom are reported in Table S7.

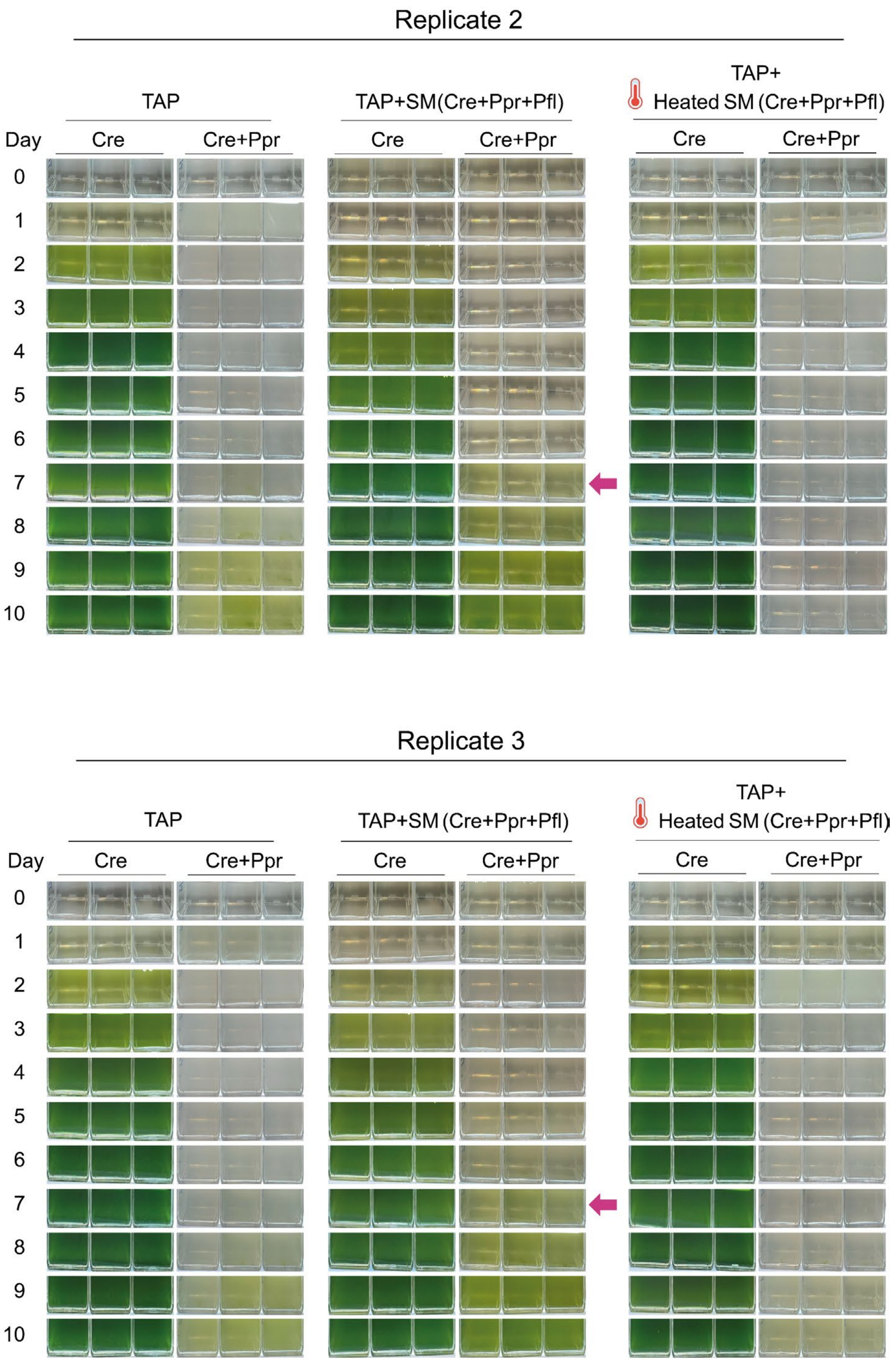


**Fig. S8 | A heat-sensitive factor(s) from *P. fluorescens* contributes to protegencin inactivation.** Cocultures of *C. reinhardtii* (Cre) and *P. protegens* (Ppr) in non-heated and heat-treated spent medium from the tripartite culture [SM (Cre+Ppr+Pfl)]. The magenta arrow indicates Cre recovery. Independent biological replicates 2 and 3 are shown. Independent biological replicate 1 is present in Fig. 6A.

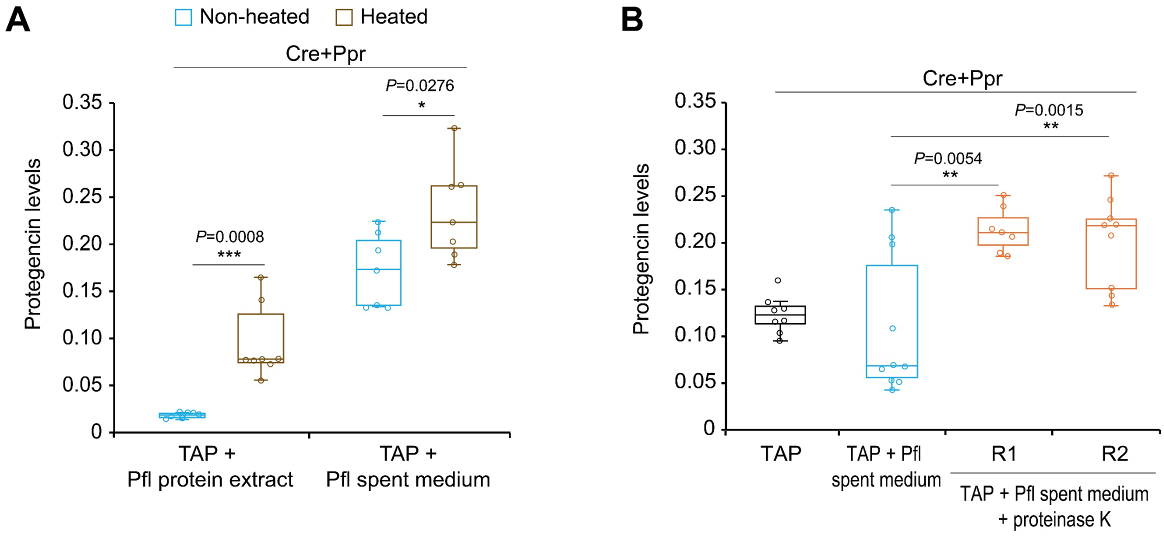


**Fig. S9 | Reduction of protegencin leves by enzyme(s) from *P. fluorescens*. A** Protegencin levels in Cre+Ppr cocultures grown in non-heated or heat-treated Pfl spent medium or protein extract. In the Pfl protein extract, 7 independent cells were measured under non-heat-treated conditions and 8 under heat-treated conditions. In the Pfl spent medium, 7 independent cells were measured under each condition. The experiment was independently repeated; the Cre monoculture control was omitted in this replicate. **B** Protegencin levels in Cre+Ppr cocultures in the spent medium of Pfl treated with proteinase K. Proteinase K concentration was 50 μg $ml^{-1}$. Two independent datasets (R1 and R2) are shown. In TAP medium, 8 independent cells were measured. In *P. fluorescens* spent medium, 10 independent cells were measured. For *P. fluorescens* spent medium treated with proteinase K, 9 and 8 independent cells were measured in the two independent replicates, respectively. **A-B** Cre: *C. reinhardtii*, Ppr: *P. protegens*, Pfl: *P. fluorescens*. Protegencin levels were quantified by Raman microspectroscopy using the ratio of the triple-bond signal at (2170 ± 100) $cm^{-1}$ to the reference signal at (1635 ± 80) $cm^{-1}$. The data are presented as box-and-whisker plots, with the center line indicating the median, the box representing the interquartile range (IQR), and the whiskers extending to the minimum and maximum values. Individual values are shown as scattered points, including both inner and outlier points. Statistical significance was determined using an unpaired two-tailed Welch's *t*-test (unequal variances). Significance levels are indicated as follows: $^*P < 0.05$, $^{**}P < 0.01$, $^{***}P < 0.001$. Exact *P*-values, *t*-values and degrees of freedom are reported in Table S8C, D, respectively.
